# Connectivity-based Meta-Bands: A new approach for automatic frequency band identification in connectivity analyses

**DOI:** 10.1101/2023.03.30.534879

**Authors:** Víctor Rodríguez-González, Pablo Núñez, Carlos Gómez, Yoshihito Shigihara, Hideyuki Hoshi, Tola-Arribas Miguel Ángel, Mónica Cano, Guerrero Ángel, David García-Azorín, Roberto Hornero, Jesús Poza

## Abstract

The majority of electroencephalographic (EEG) and magnetoencephalographic (MEG) studies filter and analyse neural signals in specific frequency ranges, known as “canonical” frequency bands. However, this segmentation, is not exempt from limitations, mainly due to the lack of adaptation to the neural idiosyncrasies of each individual. In this study, we introduce a new data-driven method to automatically identify frequency ranges based on the topological similarity of the frequency-dependent functional neural network. The resting-state neural activity of 195 cognitively healthy subjects from three different databases (MEG: 123 subjects; EEG_1_: 27 subjects; EEG_2_: 45 subjects) was analysed. In a first step, MEG and EEG signals were filtered with a narrow-band filter bank (1 Hz bandwidth) from 1 to 70 Hz with a 0.5 Hz step. Next, the connectivity in each of these filtered signals was estimated using the orthogonalized version of the amplitude envelope correlation to obtain the frequency-dependent functional neural network. Finally, a community detection algorithm was used to identify communities in the frequency domain showing a similar network topology. We have called this approach the “Connectivity-based Meta-Bands” (CMB) algorithm. Additionally, two types of synthetic signals were used to configure the hyper-parameters of the CMB algorithm. We observed that the classical approaches to band segmentation reflect the underlying network topologies at group level for the MEG signals, but they fail to adapt to the individual differentiating patterns revealed by our methodology. On the other hand, the sensitivity of EEG signals to reflect this underlying frequency-dependent network structure is limited. To the best of our knowledge, this is the first study that proposes an unsupervised band segmentation method based on the topological similarity of functional neural network across frequencies. This methodology fully accounts for subject-specific patterns, providing more robust and personalized analyses, and paving the way for new studies focused on exploring the frequency-dependent structure of brain connectivity.

## 1. Introduction

The brain is the most complex biological system of the human body [1]. It is composed by a myriad of neurons continuously communicating by synapses. There are many neuroimaging techniques that allow the acquisition of the electromagnetic activity generated by these synapses, such as electroencephalography (EEG) and magnetoencephalography (MEG). These neurophysiological techniques have a high temporal resolution (in the order of milliseconds). This enables the capture of faster brain activity, which is of great interest in a dynamic system as the brain [2]. EEG and MEG (M/EEG) also have relevant differences between them: while EEG has a reduced cost, portability, and is more widespread in clinical settings [3], MEG is contactless, and more robust against volume conduction effects [4, 5]. Furthermore, although both techniques record the same sources, EEG is sensitive to radial and tangential sources, whereas MEG is only sensitive to the latter ones [6]. It has been proposed to combine both acquisition techniques to optimize the information obtained and increase the spatial resolution of the recordings [4, 7, 8].

There are many ways of analysing M/EEG signals. One such way is by individually assessing the time course of each sensor or brain region. In this way, we can obtain information about the behaviour of individual neuronal groups, what is known as “local activation” analyses [9]. Nonetheless, by analysing only the local patterns we are losing lots of information. It is noteworthy that neurons interact between them, forming dynamical networks that emerge via chemical and electrical processes [10, 11]. Different brain regions are involved in these networks, whose interactions support higher cognitive functions, such as deduction or reasoning [12]. There is a huge amount of studies devoted to assess these interactions [13], which are known as “functional connectivity” (FC) analyses [14].

It is noteworthy that FC studies typically analyse the neural interactions in specific frequency ranges: the well-known “canonical” frequency bands (*i.e.*, delta, theta, alpha, beta-1, beta-2, and gamma). These bands are supported by abundant literature, and they reflect certain brain patterns with extensively proved physiological meaning [15, 16]. Moreover, as canonical frequency bands are widespread, they provide a common framework that ease the comparisons across subjects and studies. However, there are several issues related with their definition: i) they were specified about 80 years ago, when the techniques for the acquisition of neural activity were remarkably different than those currently available [17, 18, 19, 20, 21]; ii) there is a noticeable inconsistency in their definition, with different band boundaries across studies [22]; iii) most of the current analysis methods, such as FC, had not been developed when the canonical frequency bands were introduced and, as a consequence, they were defined based on local activation patterns, thus losing relevant features of the neural signals; and iv) the canonical frequency bands do not account for individual idiosyncrasies at subject-level. Other approaches to frequency band segmentation tried to address the latter issue, proposing subject-adaptive frequency bands [23, 24]. These methods assess the distribution of the spectral content of the signals to specify the personalized frequency band boundaries and, consequently, they also rely on local activation patterns [23, 24]. In this way, they take into account some of the individual idiosyncrasies, but a great level of homogenization is still present, as they use fixed parameters such as the number of bands or their bandwidth. FC patterns reflect the global interaction between brain regions, but also the local synchronization, thus summarizing a richer variety of neural properties [25, 26]. Hence, it is reasonable to consider whether an alternative, automatic, FC-based, and subject-specific frequency band segmentation can be appropriate.

As we have mentioned, the majority of FC studies using M/EEG employ frequency bands with fixed frequency ranges. This could be a confounding factor, as it can be hiding relevant results by grouping frequencies with very different connectivity features [22]. In this paper, we propose, to the best of our knowledge, the first methodology that allows to find unsupervised data-driven frequency bands based on the connectivity topology. The methodology proposed here automatically groups frequency bins based on the topological similarity between them. We have called this new methodology the “Connectivity-based Meta-Bands” (CMB) algorithm. Throughout this paper, the term meta-band is used to name the frequency ranges that our methodology identifies as belonging to the same community based on their network topology. The CMB algorithm allows to perform connectivity analyses accounting for the underlying frequency structure, thus having personalized analyses that fully adapt to the individual idiosyncrasies of neural activity of each subject.

Based on the hypothesis that we can automatically define user-specific frequency ranges to be used in connectivity analyses, the following research questions are posed in the manuscript: i) how is the topology of the underlying frequency-dependent neural network structure? and ii) can we define adaptive frequency bands that are a better fit for this network structure than canonical frequency bands? To answer these questions, the main objective of this paper is to develop a FC-based, unsupervised, and subject-specific frequency band segmentation approach. This would allow to perform subsequent analyses without the bias factor of considering the frequency bands to be fixed across subjects.

## 2. Materials

### 2.1. Synthetic signals

In order to assess the performance of the methodology under controlled conditions, it was tested using synthetic signals with known ground-truth. Two different types of signals were used: i) amplitude-coupled synthetic signals, and ii) M/EEG-like synthetic signals.

#### 2.1.1. Amplitude-coupled synthetic signals

These signals were designed to gain a better understanding of the performance of the methodology, as well as to define its limitations while keeping the computational burden in an acceptable level. Three minutes of activity in 20 channels were generated with a sampling frequency of 1000 Hz, and segmented in 5-second epochs. As the signals were stationary, we considered that 3 minutes of activity was enough for a proper operation of the CMB algorithm. These signals were used to assess the influence of different hyper-parameters on the results, which will be defined below: filter order, sampling frequency, bandwidth, band overlapping, and frequency resolution.

These signals were generated by means of an amplitude modulation (AM) in specific frequency ranges and channels (that determined our underlying meta-bands), and a low level of white Gaussian noise in the broadband (−19 dBW). The final signal, *s*(*t*), in each sensor *i* was constructed as follows:

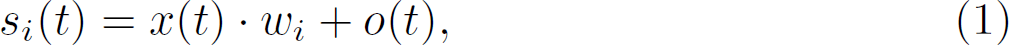

where *x*(*t*) is the modulated signal (equation 2), *w_i_* the binary weight used to construct the underlying connectivity matrix for each sensor *i*, and *o*(*t*) a white Gaussian noise signal. In this way, *x*(*t*) is the same for all the sensors, while the coupling is determined by *w_i_*, with “1” indicating coupling, and “0” not coupling. The AM signal *x*(*t*) is defined as follows:

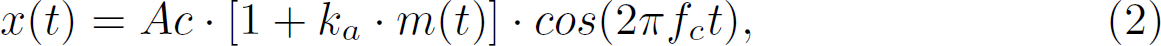

where *A_c_* is the carrier amplitude, *k_a_* the modulation index, *m*(*t*) the modulating signal, and *f_c_* the carrier frequency. Finally, the modulating signal *m*(*t*) is defined as:

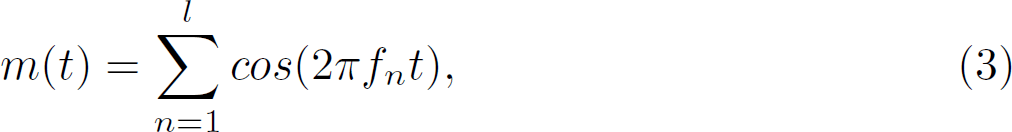

with *l* being the number of cosines composing the signal, and *f_n_* the frequency of each cosine. Thus, by varying *f_c_*, *f_n_* and *l*, we can obtain signals with the AM in different frequencies and with different bandwidths. In the simulated scenarios where more than two AM are required, the two signals are summed, *i.e. s_i_*(*t*) = *s_i_*_1_(*t*) + *s_i_*_2_(*t*). Further details on the generation of these signals can be found in section 1.1 of the Supplementary Material.

#### 2.1.2. M/EEG-like synthetic signals

To evaluate the performance of the CMB algorithm with signals closer to brain physiological time courses, M/EEG-like synthetic signals were generated. These signals contain a similar data distribution to that of real M/EEG recordings, thus providing a more realistic scenario. Nonetheless, they have an increased computational burden compared to the amplitude-coupled synthetic signals. These signals have the same number of neural sources as the original M/EEG signals (*i.e*, 68 neural sources), the same duration in time, and are also divided into 5-second epochs.

We employed 5 real source-level MEG recordings as the basis to create these signals. First of all, we took advantage of surrogate data testing to eliminate the static FC interactions from the 5 recordings [27, 28]. Specifically, we used the amplitude adjusted Fourier transform (AAFT), which is an improved version of the phase randomization method of surrogate data construction which preserves the amplitude distribution of the data [29, 30]. This algorithm is capable of destroying the static FC of a time series while preserving the amplitude distribution and the linear structure of the data [27, 28]. That is, after applying AAFT, we have 5 synthetic M/EEG-like signals similar to the original ones, but with a negligible degree of static FC between them. Then, the coupling between these signals (*i.e,* the FC) was generated by means of filtering and weighted summing processes in specific frequency ranges. The resulting signals have non-negligible FC only in the defined frequency ranges.

These synthetic signals were used to gain deeper insights on how the sampling frequency affects the meta-bands. In addition, we employed these signals to find out whether our methodology is capable of recovering, at least, 6 underlying bands with and without a gap with negligible FC between them (see section 3.3). More details about the generation of these signals can be found in section 1.2 of the Supplementary Material.

### 2.2. Real M/EEG recordings

The CMB algorithm was also applied to analyse neural activity of a cohort of 195 subjects from three different databases: 123 from an MEG database, and 27 and 45 for two independent EEG databases. For all the databases, 5 minutes of resting-state eyes-closed brain activity were recorded. All patients were asked to remain awake and still during the recordings. The neural activity was monitored in real time to prevent subjects from falling asleep [11, 31]. All the subjects included in the study had no history of neurological or psychiatric disorders. In addition, they did not take medication that could have an influence on M/EEG activity. Although the CMB algorithm can also be applied to task-related paradigms, their dynamical characteristics require further considerations on the CMB algorithm that should be addressed in future studies, as they increase the complexity of the required tests. Thus, we decided to employ the simplest paradigm, based on considering resting-state recordings, to provide the initial evaluation of the method. Furthermore, the resting-state paradigm is widely used by the scientific community to study intrinsic brain behaviour and neural alterations associated with different pathologies [32, 33, 22]. Hence, this paradigm is useful in clinical setting in which other protocols are difficult to apply (or even impossible), for example with patients suffering from dementia.

#### 2.2.1. Characteristics of M/EEG recordings

MEG activity was recorded from 123 cognitively healthy controls using a 160-channel axial gradiometer MEG system (MEG Vision 60C, Yokogawa Electric, sampling rate 1000 Hz). The sample is composed of 61 females and 62 males with an age of 48.3*±*16.1 years (mean*±*standard deviation, SD).

The first EEG database (EEG_1_) was comprised by 27 healthy females. Neural signals were recorded with a 32-channel EEG system (actiChamp Plus, BrainVision) at the Clinical University Hospital, Valladolid (Spain) and at the Institute for Research & Innovation in Health of the Porto University (Portugal). The electrodes were placed according to the 10-20 system using a common reference: C3, C4, Cz, CP1, CP4, CP5, CP6, F3, F4, F7 F8, Fz, FC1, FC2, FC5, FC6, Fp1, Fp2, FT9, FT10, O1, O2, Oz, P3, P4, P7, P8, Pz, T7, T8, TP9, TP10. The age of the sample was 30.8*±*6.6 years (mean*±*SD).

The EEG activity of the second EEG database (EEG_2_) was recorded from 45 cognitively healthy elderly controls. Brain activity was recorded with a 19-channel EEG system (XLTEK^®^, Natus Medical) at the Department of Clinical Neurophysiology of the “Río Hortega” University Hospital, Valladolid, Spain. Electrodes were placed according to the international 10-20 system using a common average reference: Fp1, Fp2, Fz, F3, F4, F7, F8, Cz, C3, C4, T3, T4, T5, T6, Pz, P3, P4, O1, and O2. The sample was composed by 31 females and 14 males, with an age of 76.3*±*4.0 years (mean*±*SD).

All the studies were conducted according to the Code of Ethics of the World Medical Association (Declaration of Helsinki). The Ethics Committee of the Hokuto Hospital (Obihiro, Japan) approved the study for the MEG database (code #1001 and #1038); the Ethics Comitee of the Clinical University Hospital (Valladolid, Spain) and Institute for Research & Innovation in Health (Porto, Portugal) approved the study for the EEG_1_ database (codes PI19-1531 and 5/CECRI/2020); and the Ethics Comitee of the “Río Hortega” University Hospital (Valladolid, Spain) approved the study for the EEG_2_ database (code 36/2014/02). All participants and caregivers were informed about the research and study protocol and gave their written informed consent.

#### 2.2.2. Preprocessing of the M/EEG recordings

A similar preprocessing pipeline was conducted for the three databases [34]: i) filtering the signals with two finite impulse response filters: a band-pass filter to limit noise bandwidth (1-70 Hz) and a band-stop filter (49-51 Hz) to remove powerline noise; ii) independent component analysis to remove artifacted components, by means of the Infomax algorithm, to visually identify and remove components related with artifacts, and iii) visual selection of 5-second artifact-free epochs. For the MEG database 5.2 ± 3.2 (mean ± standard deviation) ICA components were rejected and 9.0 ± 6.3 epochs were visually discarded. Also, for the EEG_1_ database 6.7 ± 3.7 ICA components were rejected and 13.9 ± 8.1 epochs were visually discarded. Finally, for the EEG_2_ database 3.0 ± 2.0 ICA components were rejected and 14.4 ± 7.4 epochs were visually discarded.

Additionally, the increased number of sensors of the MEG recordings allowed the application of an additional artifact rejection step (the SOUND algorithm [31, 35]) before the filtering. This algorithm was applied using an anatomical template and a *λ*_0_ = 0.1, according to the values suggested in a previous study [31].

#### 2.2.3. Source reconstruction

MEG and EEG time series were reconstructed at source-level using the Weighted Minimum Norm Estimation (wMNE) method [36]. This algorithm restricts the sources of the inverse problem by minimizing the energy (*l*_2_ norm) while weighting deep sources to facilitate their detection [36]. This algorithm was selected as it is widely used in the context of MEG and EEG source localization [34, 37, 38, 39]. Of note, it can be argued that a more appropriate source inversion algorithm could be selected for a specific signal modality (as example, sLORETA for EEG [40]). However, in this study the source inversion process is not intended to unveil the real underlying sources, but to establish a common space for all three databases, while keeping their processing pipelines as similar as possible. The wMNE implementation is freely available in Brainstorm toolbox (http://neuroimage.usc.edu/brainstorm) [41].

An anatomical forward model was created using the ICBM152 template [42, 43]. Based on it, a three-layer (brain, skull, and scalp) realistic head model was generated with a boundary element method using the OpenMEEG software [34, 44]. The source space was limited to the cortex using a total number of 15000 sources [34]. The sources were restricted to be normal to the cortex [34]. Finally, after flipping the sources of opposite direction to avoid blurring of neighbouring generators [45], the 15000 sources were grouped in the 68 regions of interest (ROIs) according to the Desikan-Killiany atlas [34, 46, 47].

## 3. Methods: Connectivity-based Meta-Bands

### 3.1. Computation of the functional networks

Preprocessed source-level time signals were used to estimate the functional networks. In Figure 1 the analysis steps of the study are illustrated. Further details on the filtering process, and the estimation of the connectivity are included in subsequent sections. The steps carried out to build them are summarized below.

(i). Define the algorithm parameters (filter order, sampling frequency, frequency resolution, filter boundaries, etc.).
(ii). Z-score transform the signals.
(iii). Filter the signals with a narrow bandwidth.
(iv). Orthogonalize the signals.
(v). Estimate the connectivity by means of the amplitude envelope correlation (AEC).
(vi). Repeat steps 3-5 for each of the frequency ranges defined in step 1. The resulting matrices have the following dimensions: *N × M × M × Q*, where *N* is the number of frequency bins, *M* the number of sources, and *Q* the number of epochs.
(vii). Extract communities across the frequency dimension by means of the Louvain GJA algorithm. These frequency-dependent communities are what we refer to as meta-bands.

**Figure 1:**
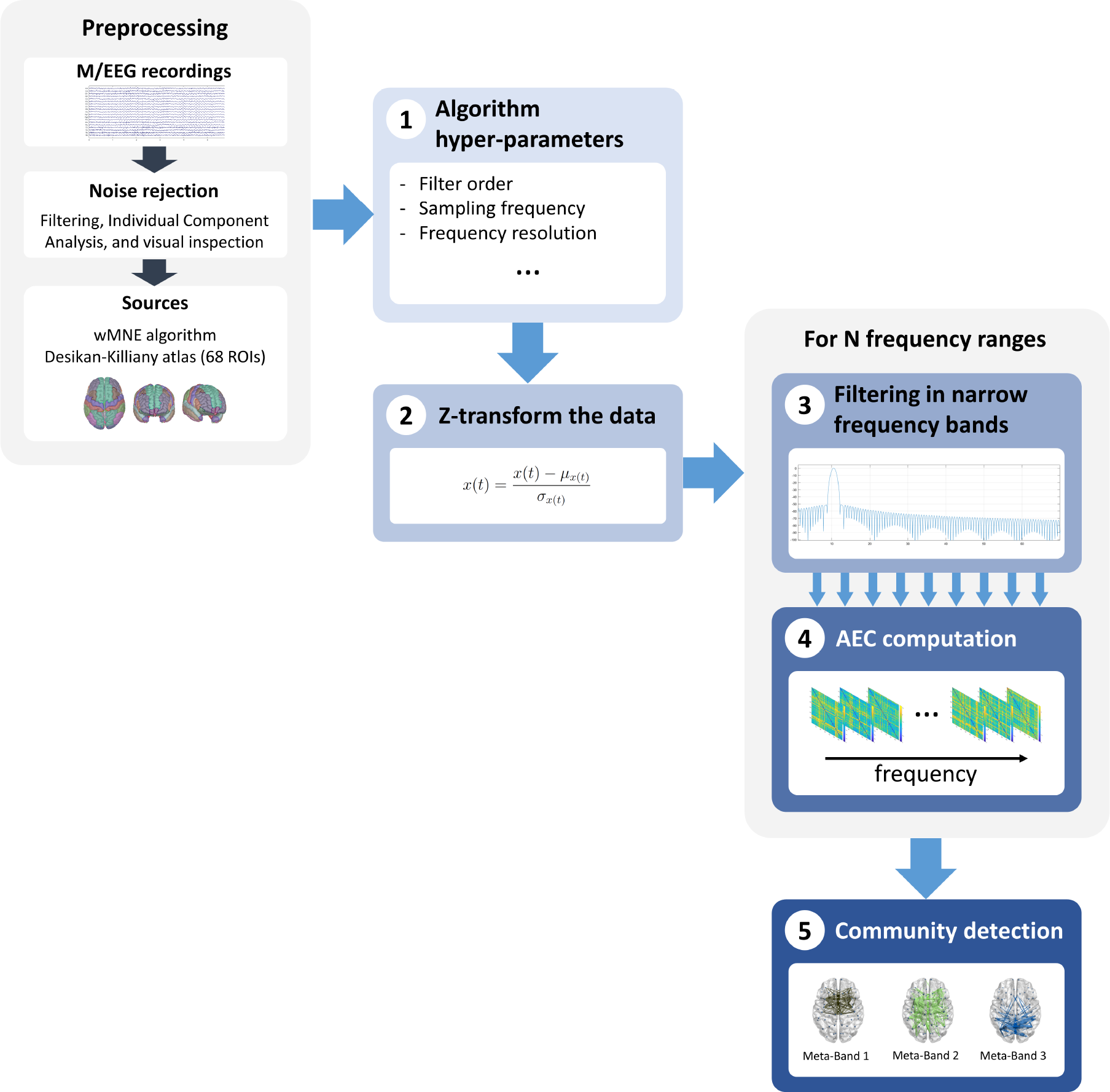
Flow diagram of the study. **(1)** Definition of the algorithm hyper-parameters based on the characteristics of our study. **(2)** Normalization of the data by means of a Z-transformation. **(3)** Filtering of the signal in narrow (1 Hz) bands. **(4)** Computation of the frequency-dependent FC by means of the amplitude envelope correlation in its orthogonalized version. Steps **(3)** and **(4)** are repeated for each frequency range defined in step **(1)**. **(5)** Community detection to estimate the meta-bands by means of the Louvain GJA algorithm.

Although the operation of the CMB algorithm is considered unsupervised, some parameters were tuned during the algorithm development. To this end, we took advantage of the synthetic signals previously described (both amplitude-coupled and M/EEG-like). Of note, although the algorithm displays satisfactory performance with the default parameters, they can be modified depending on the user requirements, such as lower computational burden or different frequency range of interest.

#### 3.1.1. Filtering process

A narrow-band filter bank was employed to filter the signals, employing a Hamming window to minimize the filter ripple [48]. The MATLAB *filtfilt* function was used to achieve a filtering process with no phase distortion [48]. Furthermore, to avoid the edge effect of the filter, the filtering process was performed considering the whole recording (before the segmentation into 5-s epochs); in addition, the first and last 5-s epochs of the synthetic and real signals were discarded to avoid the influence of filter transient effects in the results [48]. Different filter orders were assessed to evaluate the algorithm performance (namely 100, 250, 500, 750, 1000, 1250, 1500, 2000, 2500, and 3000). Additionally, the filtering process was carried out with a 50% overlap between filters to increase the frequency resolution and smooth the evolution of the frequencydependent networks.

#### 3.1.2. Connectivity estimation: orthogonalized Amplitude Envelope Correlation

The selection of the connectivity metric is a decision of utmost importance for the application of the CMB algorithm, as different connectivity metrics are likely sensitive to different aspects of brain dynamics [49, 50]. Several reasons led us to consider the orthogonalized version of the AEC. Firstly, thanks to the orthogonalization process, this metric is robust against volume conduction and field spread effects, which are major confounding factors in connectivity analyses [14]. Additionally, it has been proven to be robust and consistent, displaying a high level of reproducibility and repeatability; these features have been previously considered as quality markers for connectivity metrics [14, 50]. Also, as the mathematical formulation of the AEC is simple, it is easier to interpret than other more mathematically convoluted parameters [51, 14]. Furthermore, it is sensitive to the alterations of neurological disorders in functional connectivity, which is relevant for future potential clinical implementations of the CMB algorithm [50]. Finally, amplitude-based metrics have been adapted to compute instantaneous connectivity patterns, which have been successfully used to identify brain states from electrophysiological signals [11, 52, 53].

The AEC is an amplitude-based connectivity metric [51]. To calculate the AEC, the power envelope of the signals is computed as the absolute value of the Hilbert transform. Finally, the AEC is calculated as the Pearson correlation between the two signal envelopes. This metric was used due to its robustness, simplicity, and consistency across studies [14, 50]. Before computing the AEC, a pairwise orthogonalization was applied to the signals in order to avoid spurious correlations due to volume conduction and field spread effects [54]. The AEC is used to obtain the frequency-dependent connectivity matrices of the filtered signals from the previous step. Before performing the community detection, the connectivity matrices of all the trials of each subject were averaged. Otherwise, the high dimensionality of the data would make the community detection impossible due to the computational burden.

### 3.2. Automatic frequency band identification

#### 3.2.1. Recurrence plots

Recurrence plots (RP) are two-dimensional matrices that ease the detection of clusters and patterns of periodicity in the data they represent [55, 56]. Originally, they were designed to detect time patterns in dynamical systems [55]. They are formulated as follows [56]:

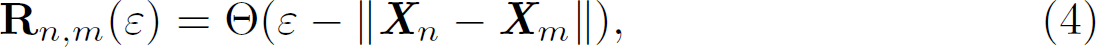

where **X***_n_* is the time course at time *n*, *ε* is a threshold, Θ(·) the Heavyside function, and ‖ · ‖ the norm. Nonetheless, in this study, we replaced the norm by the Spearman correlation between AEC matrices to avoid the selection of an arbitrary threshold [53]. Furthermore, as we are not working in the time domain but in the frequency domain, the RPs were defined as follows [11, 53]:

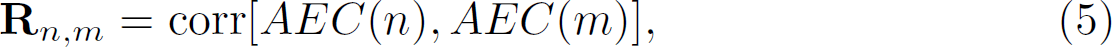

where *AEC*(*n*) and *AEC*(*m*) are the connectivity matrices for the frequencies *n* and *m*, respectively. Thus, the RP is used to identify frequency-dependent clusters that group connectivity matrices with a similar topological structure.

#### 3.2.2. Louvain community detection algorithm

Community detection algorithms have been used to detect clusters of brain regions [57, 58], or temporal brain meta-states [11, 53, 59]. Here, this methodology was used to detect repeated network patterns along the frequency dimension that can be identified as meta-bands. As we wanted to implement the frequency band identification in an unsupervised fashion, the Louvain GJA algorithm was used [11]. This method does not require the *a-priori* definition of the number of communities to discover, overcoming, in this sense, other algorithms such as *k* -means clustering, or non-negative tensor factorization [11, 13, 53, 60, 61].

For this task, the RP constructed in the previous section is considered as a graph, with each of the frequency-dependent connectivity matrices being the nodes, and the Spearman correlation between them being the edges. Communities were then discovered in the RP by means of the Louvain GJA method [62, 63]. The Louvain GJA method is an iterative algorithm that maximizes the modularity of its solution by assigning each node to its own community and then finding modularity maxima by moving nodes to other communities [58]. This method has been proven to perform well with poorly-defined communities [58]. As the algorithm is non-deterministic, it was run 250 times, selecting the solution with highest modularity [11].

To ease the comparison across subjects, we conducted a per-group community detection pipeline [52]. To do so, RPs were constructed by concatenating all the frequency-dependent connectivity matrices of every subject in a database [11]. Hence, community detection was carried out on square connectivity matrices with dimension *N*, where *N* is the number of subjects in a database multiplied by the number of frequency bins. The network topologies of the meta-bands were obtained by averaging all the connectivity matrices assigned to each meta-band [11].

#### 3.2.3. Frequency-band Activation Sequence (FAS), Attraction Strength (AS), and Degree of Dominance (DoD)

A metric was defined based on the community extraction performed by the Louvain GJA method, the frequency-band activation sequence (FAS). This is a symbolic function that contains the information about the dominant metaband network topology for each frequency bin (*i.e.*, the functional connectivity configuration assigned by the Louvain algorithm to that bin). Metrics containing information about changes between meta-states have been previously used to characterize brain state transitions [11, 60, 64]. Nonetheless, the FAS only contains information regarding to which meta-band belongs each frequencydependent connectivity matrix at each frequency bin. Consequently, in order to complement this information, two new metrics were defined: the attraction strength (AS) and the degree of dominance (DoD). These parameters measure how well the network topology of a specific frequency fits its assigned meta-band.

The AS is the Spearman correlation between each frequency-dependent connectivity matrix and the topology of the dominant meta-band (*i.e.*, the meta-band to which it has been assigned). It is calculated as follows:

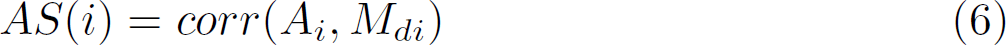

where *corr*(·) is the Spearman correlation, *M_di_* the network topology of the dominant meta-band in the frequency *i*, and *A_i_* the connectivity matrix in the frequency *i*.

The DoD is defined as the correlation between each connectivity matrix and the dominant meta-band, minus the average correlation between that connectivity matrix and the non-dominant meta-bands. It is calculated as follows:

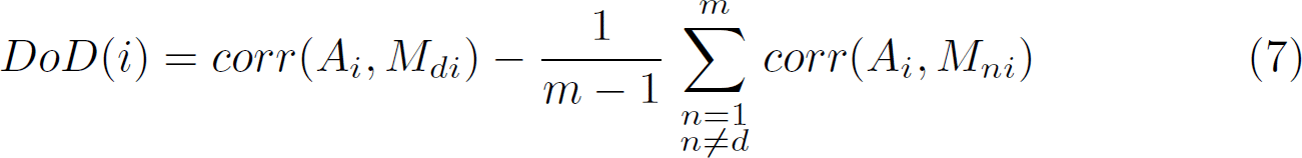

where *M_ni_* is the network topology of each of the non-dominant meta-bands in the frequency *i*. By analysing the complementary information obtained from these metrics (FAS, AS, and DoD), we can deepen our understanding of the frequency structure of brain FC.

### 3.3. Assessing the algorithm performance by means of synthetic signals with known ground-truth

The performance of the CMB algorithm was evaluated with the two types of synthetic signals (amplitude-coupled and M/EEG-like). Two different types of tests were performed: i) evaluation of the algorithm to assess whether it is able to detect meta-bands in hypothetical demanding situations; ii) tuning of the parameters to assess their impact on the results.

In the first scenario, the following parameters were evaluated:

- **Bandwidth:** Different values of the underlying synthetic meta-band bandwidth were tested to find out whether there is a hard limit where the algorithm is unable to recover the underlying meta-band. This parameter was tested for amplitude-coupled synthetic signals.
- **Band overlap:** The presence of close frequency bands (or even overlapping ones) could also be a problem for the algorithm. This parameter was tested for amplitude-coupled synthetic signals.
- **Number of meta-bands:** This simulation was performed to assess whether the number of underlying meta-bands that the CMB algorithm is capable of recovering. This parameter was tested for M/EEG-like synthetic signals.
- **Continuous bands:** The previous test of the capability of the algorithm of recovering multiple bands was repeated again with two different band configurations. First, we considered a scenario where a frequency range with no underlying network structure was left between meta-bands. For example, if two meta-bands are defined in the range of interest between 1 and 70 Hz, they could range, *e.g.*, from 5 to 35 Hz and from 45 to 60 Hz, with a frequency band with no underlying defined network structure in between. Second, we also considered a scenario where the meta-bands were adjacent (*i.e.* the frequency range without structure that separated the meta-bands is no longer considered). That is, if we define two meta-bands in a frequency range of interest between 1 and 70 Hz, they could range, *e.g.*, from 1 to 40 Hz and from 40 to 70 Hz. This test was carried out to assess whether the CMB algorithm is capable to detect the boundaries not only between meta-bands but also between a meta-band and a frequency region without a specific FC pattern. This parameter was tested for M/EEG-like synthetic signals.

In the second scenario, we evaluated:

- **Filter order:** The meta-bands were generated with different filter orders to assess their impact on the results. This parameter was tested for amplitude-coupled synthetic signals.
- **Sampling frequency:** Different sampling frequencies were tested to evaluate the influence of this parameter. This parameter was tested for amplitude-coupled and M/EEG-like synthetic signals.
- **Frequency resolution:** The influence of the bandwidth of the filters of the narrowband filter bank on the meta-bands was evaluated. This parameter (and the frequency range under study) define the number of frequency bins. This parameter was tested for amplitude-coupled synthetic signals.

## 4. Results

### 4.1. Assessing the CMB algorithm performance in demanding situations

The performance of the CMB algorithm in the hypothetical demanding situations defined in section 3.3 is summarized below (a more detailed description of the results can be obtained in section 2 of the Supplementary Material):

- **Bandwidth:** The results showed wider bandwidths provide better-defined connectivity matrices, as there are more frequency bins with a given connectivity pattern. Further details are provided in section 2.1.2 of the Supplementary Material.
- **Band overlap:** The CMB algorithm is capable of detecting close, contiguous, and even overlapping meta-bands. In this latter scenario, in the frequency ranges where two meta-bands overlap, only one of them is detected. Further details are provided in section 2.1.4 of the Supplementary Material.
- **Number of meta-bands:** The CMB algorithm is able to recover any number of meta-bands. Further details are provided in section 2.2 of the Supplementary Material.
- **Continuous bands:** The CMB algorithm is able to recover adjacent and non-adjacent meta-bands. Further details are provided in sections 2.2.1 and 2.2.2 of the Supplementary Material.

### 4.2. Assessing the influence of the CMB algorithm hyper-parameters in the results

The results about the influence of the different hyper-parameters of the CMB algorithm in the meta-band detection can be summarized as follows (a more detailed description of the results can be obtained in section 2 of the Supplementary Material):

- **Filter order:** Lower filter orders yield a decreased frequency resolution to detect the underlying meta-band (*i.e.*, lower efficiency detecting the edges of the meta-bands). Also, higher filter orders lead to a noisier detection in the frequencies where no meta-band was specified and, thus, to less defined connectivity matrices with lower coupling values. In consequence, we have selected a filter order of 500 to get a good balance between the aforementioned effects. Further details are provided in section 2.1.1 of the Supplementary Material.
- **Sampling frequency:** It can be observed that lower sampling frequencies provide slightly narrower bands for a given filter order. Besides, the lower the sampling frequency, the lower regularity in the detection (*i.e.*, spurious state changes) in frequencies where no underlying band was defined. Further details are provided in sections 2.1.3 and 2.2.3 of the Supplementary Material.
- **Frequency resolution:** Higher bandwidths of the narrowband filter bank are associated with decreased frequency resolution in the results. Of note, increasing the frequency resolution increments the computational burden of the CMB algorithm. Further details are provided in section 2.1.5 of the Supplementary Material.

### 4.3. Assessing the CMB algorithm with real signals

The assessments carried out with synthetic signals helped us to decide which parameters to select when computing the CMB algorithm in real signals. First, the original sampling frequency was used for the three databases. The bandwidth of the filters was set to 1 Hz (thus being the CMB algorithm capable to detect 1 Hz bands), as lower values will dramatically increase the computational burden. As previously indicated, higher values of this factor will provoke solutions with lower frequency resolution, and less spurious meta-band transitions. This can be appreciated in Figures S20 and S21 in Supplementary Material. All the analyses were conducted between 1 and 70 Hz. Of note, the connectivity matrices around 50 Hz (47.5-52.5 Hz) were discarded as they are highly influenced by the 50 Hz power-line artifact (and its associated notch filtering process) [25]. In line with the findings observed in the synthetic signals, a filter order of 500 was employed. To ensure convergence of the Louvain GJA algorithm, the number of iterations was set to 250, although 100 iterations have been proven to be enough [11]. It is noteworthy that the specific values of these parameters can be adapted to fit the requirements of each study.

Figure 2 summarizes the meta-bands detected for the MEG database. The corresponding AS and DoD values are also depicted, as well as the topology for the three meta-bands networks detected. A fronto-medial network meta-band (brown) that is dominating approximately from 15 to 25 Hz can be observed. In addition, there is a medial network meta-band (green) that is mainly dominating in low (1-5 Hz) and high (35-70 Hz) frequencies. Finally, there is a parieto-occipital network meta-band (blue) that dominates around alpha (5-15 Hz) and beta-2 bands (25-35 Hz). Regarding the AS, it could be observed that it decreases in: i) the low frequencies; ii) the blue meta-band around alpha; and iii) from 30 Hz on. The higher AS values, the better the adjustment of individual frequency-dependent connectivity matrices with the corresponding meta-band network topology. Moreover, it can be appreciated that the DoD has two peaks: i) in the blue meta-band around alpha; and ii) in the brown meta-band. An increase of this value indicates a higher topological distance between the dominant meta-band and other meta-bands.

**Figure 2:**
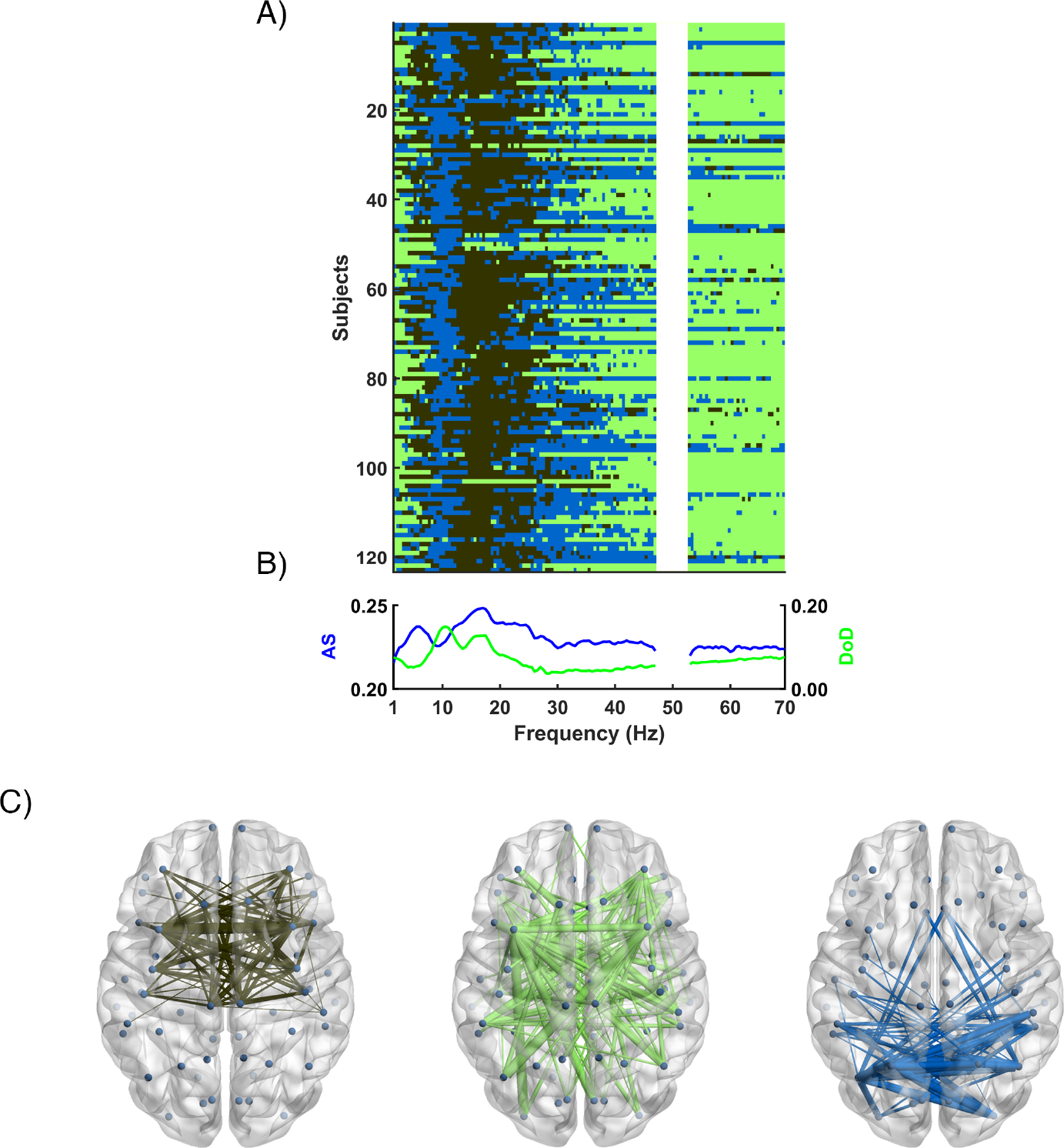
Meta-bands detected with the MEG database using a filter order of 500. A) Y-axis includes the FAS for each subject, while X-axis represents each individual frequency bin; different colours represent different communities. B) Frequency evolution of Attraction Strength (AS, blue) and Degree of Dominance (DoD, green). C) Network topologies of the three meta-bands detected; the colour of each network topology corresponds with the colour of the meta-bands in the upper plot.

Figure 3 shows the meta-bands as well as their corresponding AS, DoD, and topologies for the EEG_1_ database. The differences with the meta-bands features detected for the MEG database are noteworthy, nonetheless, it can also be observed some similarities. There is a meta-band dominating from 1 Hz to approximately 30 Hz (brown) and another meta-band dominating from 30 to 70 Hz (green). Moreover, the blue meta-band only appears in some subjects from 20 Hz on, and in a few isolated frequencies. In addition, the brown meta-band displays a parieto-occipital topology similar to the blue meta-band of the MEG database. The green meta-band displays a fronto-medial topology, and the blue one a right-medial network configuration. Finally, AS and DoD parameters show a peak around 10 Hz: while the peak of the AS is negative, the peak of the DoD is positive (similar to the behaviour of DoD in the MEG database).

**Figure 3:**
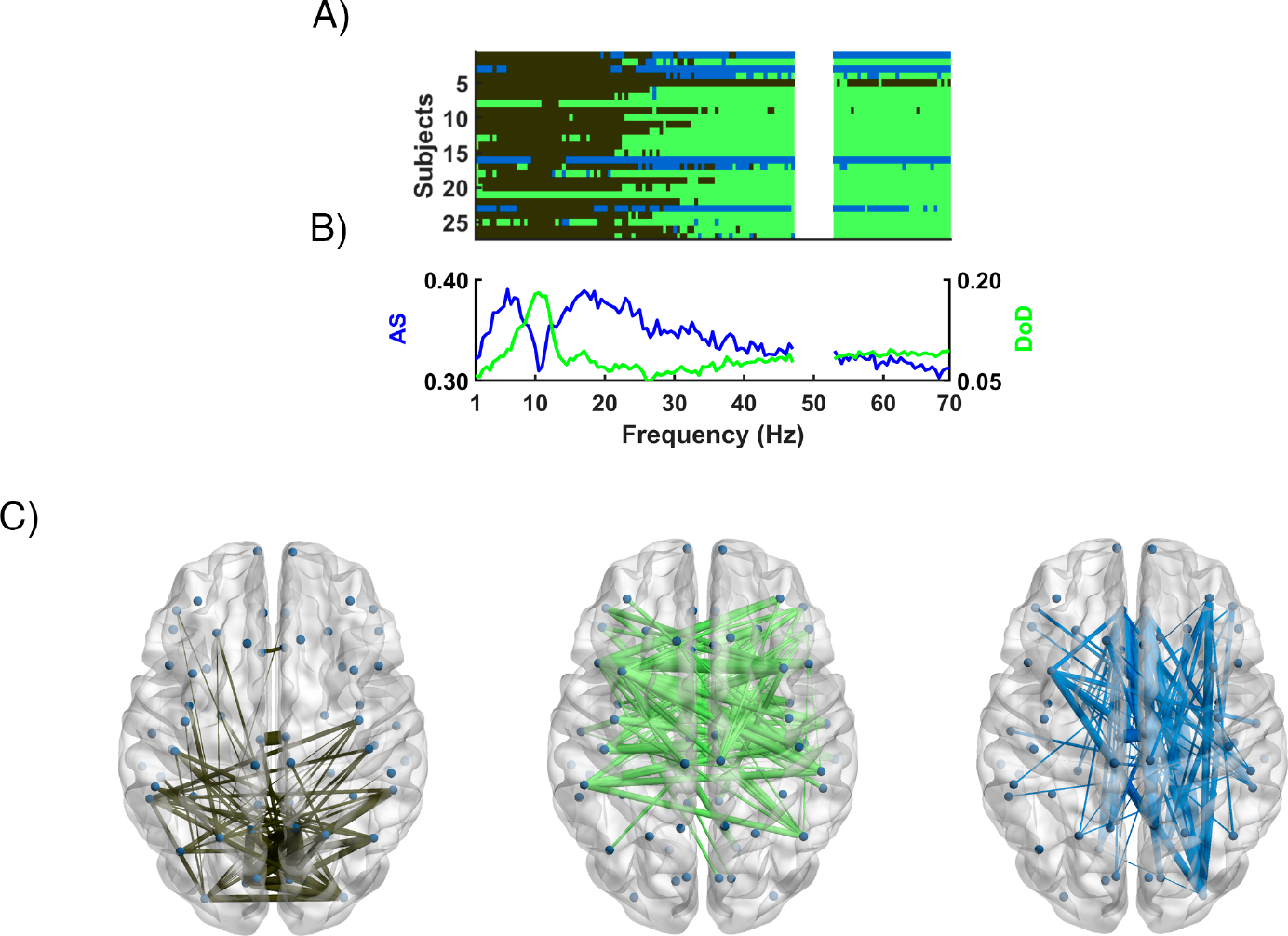
Meta-bands detected with the EEG_1_ database using a filter order of 500. A) Y-axis includes the FAS for each subject, while X-axis represents each individual frequency bin; different colours represent different communities. B) Frequency evolution of Attraction Strength (AS, blue) and Degree of Dominance (DoD, green). C) Network topologies of the three meta-bands detected; the colour of each network topology corresponds with the colour of the meta-bands in the upper plot.

Figure 4 summarizes the meta-bands as well as their corresponding AS, DoD, and topologies for the EEG_2_ database. The frequency distribution of the meta-bands is very similar to the one of the EEG_1_: the brown meta-band dominates approximately from 1 to 30 Hz; the green meta-band dominates approximately from 30 to 70 Hz; and the blue meta-band only appears in some subjects from 20 Hz on, and in a few isolated frequencies. Furthermore, the topologies of the green (fronto-medial) and blue (right-medial) meta-bands are also similar to those found in the EEG_1_ database. On the other hand, the network configuration of the brown meta-band is different from that for the EEG_1_ database, having a medial topology. Both AS and DoD present a negative peak around 10 Hz, with a steeper decline for the AS.

**Figure 4:**
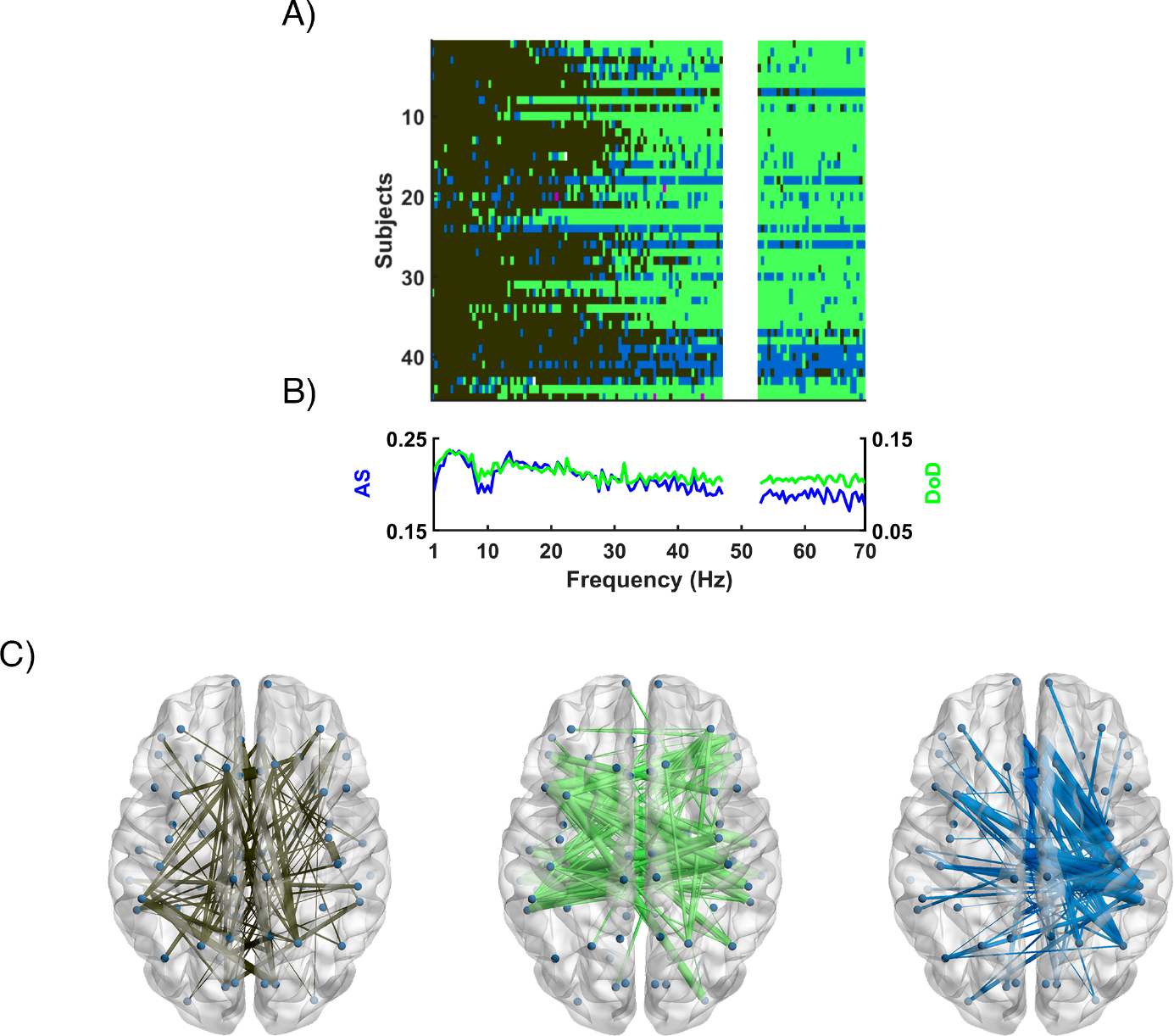
Meta-bands detected with the EEG_2_ database using a filter order of 500. A) Y-axis includes the FAS for each subject, while X-axis represents each individual frequency bin; different colours represent different communities. B) Frequency evolution of Attraction Strength (AS, blue) and Degree of Dominance (DoD, green). C) Network topologies of the three meta-bands detected; the colour of each network topology corresponds with the colour of the meta-bands in the upper plot.

The results in Figures 2, 3, and 4, but calculated with a filter order of 1500, are also included in Supplementary Material in Figures S22 (MEG database), S23 (EEG_1_ database), and S24 (EEG_2_ database). There, the noisier meta-band detection obtained when using 1500 as filter order can be appreciated. The problem of noisy detection was also observed in previous studies [11, 52].

In addition, the extent to which the CMB algorithm fits other approaches used to automatically define the frequency bands was assessed for MEG signals. To do that, we quantified the alignment of the meta-band in the alpha range (blue meta-band) with other definitions of this frequency band, as it is known that alpha band plays an important role in resting-state recordings [65]. Two definitions to estimate the alpha band were evaluated: the canonical alpha band (8-13 Hz), and an adaptive alpha band based on estimating the individual alpha frequency (IAF) and considering the alpha band as: IAF *±* 2 Hz. [66, 67]. This process was only performed for the MEG database, as the EEG databases lack a clear meta-band specific to the alpha frequencies. In Figure 5 the overlap of the detected meta-bands with the canonical (A) and adaptive (B) alpha band approaches is depicted. For the canonical approach, our alpha meta-band is present in a 59.7% of the frequency bins, while for the adaptive approach the degree of overlap of our alpha meta-band is 67.4%. In addition, in Figure 6 the modal meta-band across subjects is depicted, showing a remarkable consistency with the canonical definition of the frequency bands (shown in a bar above): delta: 1-4 Hz, theta: 4-8 Hz, alpha: 8-13 Hz, beta: 13-30 Hz, and gamma: 30-70 Hz. Besides, in that Figure the network topologies of both, the canonical bands and the meta-bands are also displayed. These network topologies support the validity of the results obtained by the CMB algorithm, as there is a remarkable coincidence between the network topology of the meta-bands and the topology of canonical bands in the frequencies where they are present: brown meta-band presents a connectivity pattern similar to those for delta and gamma bands; the network topology of the green meta-band is similar to those of theta, beta-1, and beta-2; and blue meta-band has a connectivity pattern similar to that of alpha.

**Figure 5:**
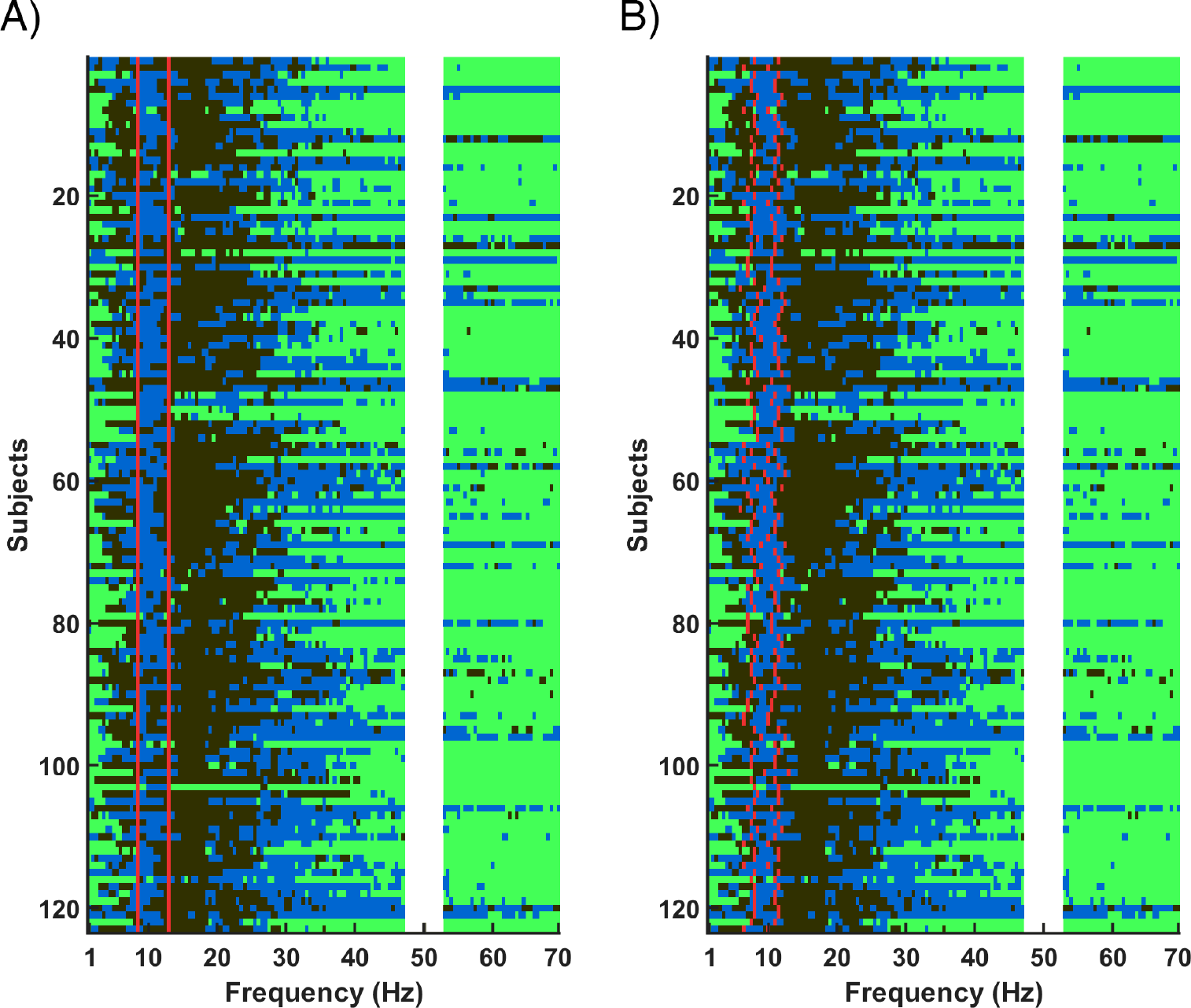
Overlapping of meta-bands (for the MEG database using a filter order of 500) with other alpha band definitions: A) canonical alpha band (8-13 Hz); B) adaptive alpha band (individual alpha frequency ± 2 Hz). The alternatives definitions of the alpha band are delimited with two red lines. The Y-axis includes the FAS for each subject, while the X-axis represents each individual frequency bin; different colours mean different communities.

**Figure 6:**
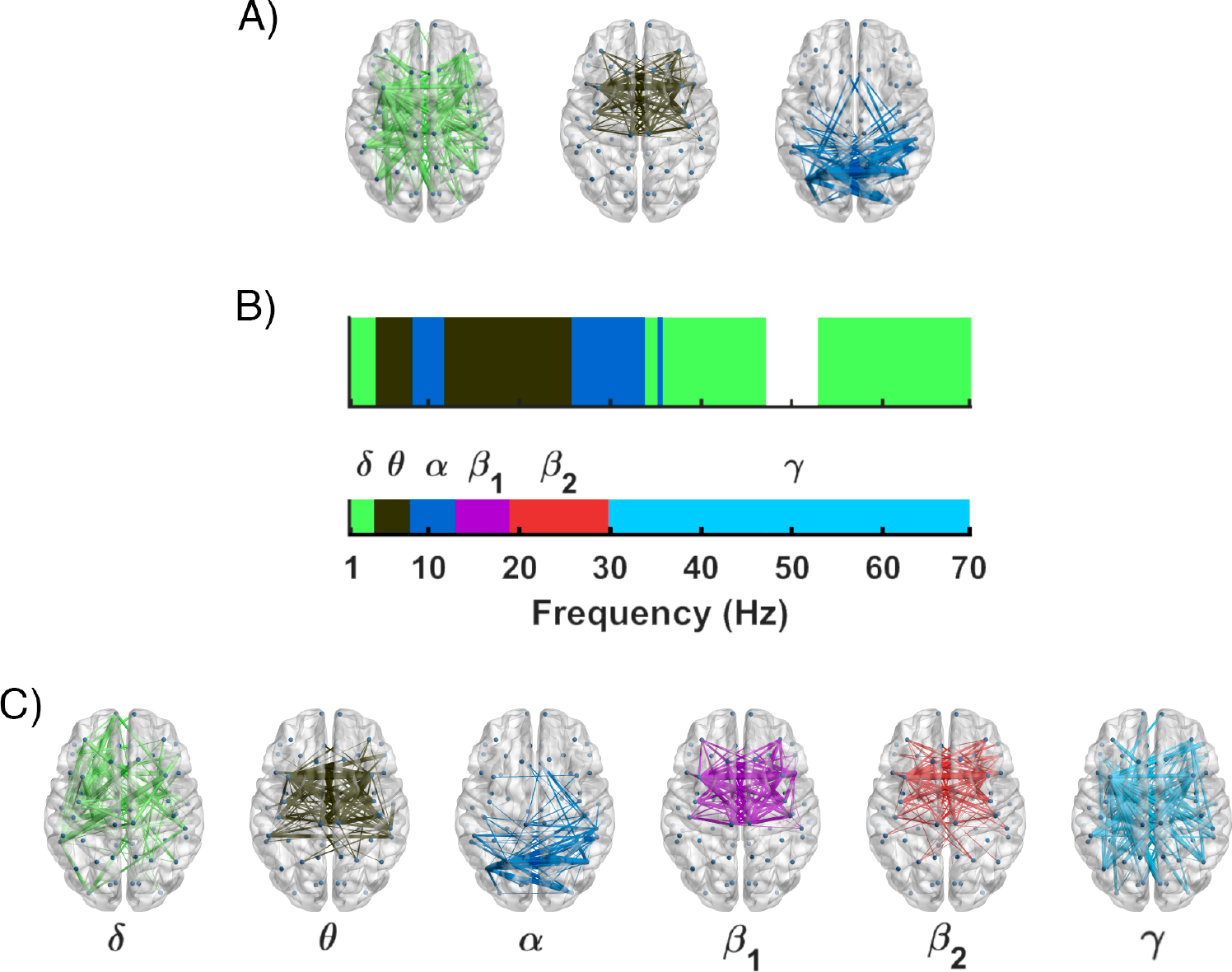
Comparison of meta-bands (for the MEG database using a filter order of 500) with canonical frequency bands. A) Network topologies of the identified metabands. B) In the upper part, the mode of the meta-bands across all the subjects is depicted, below there is a bar showing the canonical definition of the frequency bands (*δ*: 1-4 Hz; *θ*: 4-8 Hz; *α*: 8-13 Hz; *β*_1_: 13-19 Hz; *β*_2_: 19-30 Hz; and *γ*: 30-70 Hz). C) Network topologies of the canonical frequency bands. Different colours represent different communities.

## 5. Discussion

In this study, we introduced a new method to study the frequency structure of the functional connectivity in resting-state signals. We performed a community detection procedure along the frequency dimension to unveil the underlying meta-bands, which were grouped according to the similarity of their network topology. Our findings showed that: i) the CMB algorithm is sensitive to individual idiosyncrasies at subject level, providing a new framework to further understand the frequency structure of functional brain networks; ii) this frequency structure is dominated by a limited number of meta-bands (*i.e.*, connectivity topologies), that possibly play different roles depending on the frequency range; and iii) MEG is more sensitive than EEG to characterize the underlying frequency structure of functional connectivity.

### 5.1. Revisiting canonical frequency bands

The well-known canonical frequency bands are supported by a large body of studies. It is clear that they reflect underlying physiological mechanisms [15, 68]. Nonetheless, there are a variety of issues related with their definition that lead us to look for an automatic, unsupervised, and adaptive frequency segmentation in connectivity studies. The first issue is that they were defined about 80 years ago, using the rudimentary EEG systems that were available at the time [17, 19, 21]. Since then, technology and, as a consequence, recording systems have dramatically changed, and are now able to record neural activity with a highly increased quality. Because of this, it is reasonable to hypothesize that nowadays more accurate frequency bands segmentations can be performed. The next issue is related with the frequency bands limits. The literature shows a great inconsistency in the definition of the specific limits of the canonical frequency bands [22]. Thus, even though the name of the bands is preserved, their specific definition varies across studies [22]. The third issue is that analysis methods have deeply evolved since the original definition of the canonical frequency bands. The current analysis techniques are able to measure different patterns of neural activity than the ones used to define the canonical bands. For example, functional connectivity reflects global (and local) neural synchronization, while the methods used to define the canonical bands only reflect local neural synchronization [9, 25, 26]. Finally, the canonical frequency bands do not account for the individual idiosyncrasies, *i.e.*, the differentiating brain patterns across individuals. In line with this, there is an increased interest in personalized analyses, which have lead to the definition of adaptive frequency bands [23, 24]. Our methodology allows to define user-specific frequency bands, with a data-driven method that permits an increased level of personalization regarding the currently used adaptive approaches.

It has to be indicated that a direct comparison between the CMB algorithm and the canonical bands is not possible. The CMB algorithm disregards all the assumptions made by the canonical bands and, hence, the resulting frequency segmentation is so different that the direct comparison with the canonical bands is not possible: the frequency ranges are different, the number of meta-bands does not match, and the meta-bands expand across non-adjacent frequency ranges. Additionally, although the topologies of the canonical frequency bands showed a remarkable coincidence with their (approximate) corresponding meta-bands, this analysis is too simple to make direct associations between meta-bands and canonical bands. Furthermore, the limits of the canonical bands are not clear, as there is a great inconsistency in their definition [22]. Consequently, we decided to only compare the alpha band of these previous approaches with the meta-band associated with alpha (the blue meta-band) in the MEG database, as it has been observed that alpha activity dominates neural dynamics during eyesclosed rest condition [69, 70]. It can be appreciated that our blue meta-band has an occipital connectivity pattern, similar to that observed in the canonical alpha band. Of note, the adaptive alpha band showed higher coincidence (67%) with our alpha meta-band compared to the canonical alpha band (60%). This suggests that, although the traditional band segmentation approaches provide a good starting point for fitting the underlying topological structure at group-level, the individual idiosyncrasies are not fully accounted for. In this regard, the CMB algorithm is useful as it automatically adapts to the specific neural patterns of each subject, with no *a-priori* assumptions, as the canonical and adaptive approaches do.

The majority of the currently available user-specific band segmentation approaches are based on detecting spectral peaks, which can be problematic in the absence of observable peaks in the power spectral density [71]. In consequence, it is difficult to compare our meta-bands with previous studies as, to the best of our knowledge, only two previous works have assessed the frequency structure of functional connectivity [71, 72]. The study conducted by Puxeddu and colleagues [72] employed a community-detection algorithm in frequency-specific connectivity matrices. Nonetheless, the research was not aimed at proposing an alternative frequency segmentation method, but to use a multi-layer model to analyse multi-frequency EEG networks [72]. On the other hand, based on the idea that the current band segmentation approaches add uncertainty and subjectivity to the analyses, Cohen [71] proposed an alternative band segmentation approach. He used covariance matrices to identify patterns that maximally separate the spatio-temporal narrowband activity from the spatio-temporal broadband activity. Then, he grouped these patterns using the *dbscan* clustering algorithm, resulting in its user-specific bands [71]. His results showed a great variability across subjects in the number of detected frequency bands (ranging from 4 to 11), and also in the limits of the personalized alpha band (with the lower bound ranging approximately from 3 to 10 Hz, and the upper bound ranging approximately from 6 to 18 Hz). Our results support the heterogeneity in band segmentation found in this previous work, which suggests the need for individualized analyses that accounts for these user-specific idiosyncrasies. The CMB algorithm performs an individualized band segmentation derived from the connectivity-based spatio-frequency patterns obtained by means of the AEC method, as it is robust and repeatable [14, 50]. In addition, the CMB algorithm uses the Louvain GJA algorithm, as it does not *a-priori* define any parameter to achieve a satisfactory performance [11, 52]. In consequence, to the best of our knowledge, this is the first study that proposes a subject-specific frequency band segmentation based on the topological properties of the frequency-dependent connectivity matrices.

### 5.2. Unveiling the fundamental functional network configurations

Regarding the number of meta-bands detected for the different databases, a relevant conclusion can be drawn. Whereas in the MEG signals 3 clearly defined communities are observed, in the EEG databases only 2 dominant communities are detected, being the third one only present in isolated frequency bins and subjects. Of note, these results do not mean that our resting-state neural activity displays only 2 or 3 specific network topologies; they mean that there are 3 network topologies that recurrently appear across frequencies during the resting-state.

As this is the first study that extracts frequency meta-bands, it is not easy to find previous researches to which to relate the results. However, it may be interesting to consider our meta-bands as the frequential counterparts of the brain meta-states extracted on dynamic functional connectivity (dFC) analyses. The so-called brain meta-states are network patterns that are recurrently repeated over time, considering that the connectivity matrix in each time sample is a weighted combination of the identified meta-states [73, 74]. In this regard, the number of recurrent network topologies detected in the MEG database (*i.e.,* 3) is in agreement with a previous study, where 3 time-recurrent network topologies were extracted in resting-state EEG signals from healthy elderly subjects [11]. Although in the literature, different number of meta-states have been identified, its repertory is usually limited, considered to follow a heavy-tail distribution, with a few meta-stated being activated very frequently [75]. This is consider to provide a cost-efficient modular way of processing neural information [75, 76]. Thus, it is reasonable to hypothesize that the limited number of meta-band topologies that we have identified may also be a consequence of the optimization of neural information processing. In line with that, it has been proposed that one of the main variables ruling out brain organization may be economical constraints [77].

Another interesting finding is that the meta-bands are distributed across disjointed frequency ranges: for the MEG database, the green meta-band is present around the canonical delta and gamma bands, and the blue meta-band is present around canonical alpha and beta-2 bands. This contradiction could be explained by three factors. First, this can be due to a mixing effect provoked by averaging different trials from resting-state activity, which has been found to exhibit a highly dynamic repertoire of recurrent network topologies [11]. Second, it could be due to cross-frequency coupling. Several studies have addressed the potential relationships between different frequency bands, observing that these interactions play an important role in the processing and transmission of neural information [78, 79, 80]. Particularly, cross-frequency patterns have been demonstrated to play an important role in resting-state activity, as they contribute to long-range communication [81, 82]. In this regard, previous studies on this field have observed interactions between slow-frequency (delta and theta) and high-frequency (beta and gamma) oscillations [78, 79, 80], which could shape the functional network patterns at these frequency ranges. Finally, this distant coupling can be due to the fact that we are grouping frequency bands based on their underlying network topology. Hence, this does not necessarily indicate there are only 2 or 3 global meta-bands, but similar network topologies operating across different frequencies [16]. In this regard, it can be hypothesized that this is the result of the brain optimizing resources [77], having a reduced number of topologies integrating different functions depending on the frequency range they are operating [16]. It has been suggested that the brain has timescale-dependent network organizations [83, 84]. Our results extend that hypothesis indicating that some of those topologies could be repeated (at least to some extent) across frequencies. This is supported by previous studies that revealed that some graph characteristics of neural networks were preserved across frequencies [85, 86]. Besides, this can be observed in Figure 6 that compares meta-bands and canonical frequency bands in both, sequencing, and topology. This Figure shows that, although there are some bands historically considered different, from a mathematical perspective and based on the underlying network topology, they could be considered as closely related.

The blue MEG meta-band, which we have associated with the canonical alpha band, presents a parieto-occipital topology. This meta-band does not fit its corresponding frequency-dependent connectivity matrices very well; however, it does it remarkably better than the other (*i.e.*, non-dominant) meta-bands. A similar topology can be appreciated for the EEG_1_ database in its brown meta-band, which is the meta-band present around the canonical alpha band. On the other hand, the brown meta-band from EEG_2_, which is again present in the frequencies around the canonical alpha band, depicts a medial widespread topology. Around this meta-band, DoD and AS values decrease, indicating that the brown meta-band is not fitting the underlying network topology properly.

This suggests that the increased sampling frequency and spatial resolution of the EEG_1_ database have improved the sensitivity of the signals to reflect the underlying network structure, although still not to the same extent as in MEG signals. The network topology of this meta-band is similar to that observed in other studies, which found parieto-occipital networks for resting-state M/EEG signals in the alpha band [14, 87, 88]. This occipital topology could be related to the posterior areas of the default-mode network (DMN), which has been proposed to be the dominant brain activity during rest. Some studies have suggested that the DMN consist of two subnetworks: anterior DMN, and posterior DMN [89, 90, 91, 92]. This association with the posterior DMN is supported by previous studies which found that these areas were strongly active [91, 92], and highly connected [92] in the alpha frequencies during rest. Interestingly, the posterior areas of the DMN have been associated with higher cognitive functions such as the integration of information, attention, empathy, self-consciousness mental thoughts that occur during rest, and theory of mind [93, 89, 91].

The brown meta-band of the MEG signals has a fronto-central topology. Its high AS and DoD values suggests that this meta-band appropriately fits its corresponding frequency-dependent matrices, even remarkably better than the other meta-bands. A similar beta-band network topology has been observed in a previous study using MEG resting-state recordings [88]. Although this association is not as clear as for the blue meta-state, this topology could be associated with the anterior areas of the DMN [91, 92]. This network has been observed to be highly active for low frequencies (around theta), but not in beta [91, 92]. The anterior areas of the DMN are involved in high cognitive functions such as semantic integration, emotions, and decision making [94, 95]. The beta frequency activation of this meta-band may be related with the sensorimotor network, which has been reported to be very active and connected around beta frequencies, and displays a topology with some areas overlapping with the brown meta-band [91]. This network is also highly reported in intrinsic neural activity, and is involved in perception, proprioception, and motion [96, 97, 98]. On the other hand, the beta band has also been reported to assume a critical function in the formation of canonical resting-state networks [99], which implies a relevant role in the processing of information within and across cortical circuits [100], and may explain the role of brown meta-band.

### 5.3. Differences in the underlying connectivity patters between EEG and MEG

Of note, it was observed that a similar wide-sense frequency structure is detected in MEG and EEG. However, the results obtained from the EEG databases are missing a specific meta-band around the canonical alpha band that the CMB algorithm is able to recover in the MEG database. There are several differences between the MEG and EEG datasets that could explain the decreased EEG sensitivity to the underlying frequency structure: reduced signal-to-noise ratio [101, 102], decreased robustness against volume conduction effects [4], reduced sampling frequency, lower number of sensors, or reduced sample size, among others. To assess the latter factor, we performed an additional test by recalculating the meta-bands detected for the MEG database but only including in the analysis 27 random subjects (as in the EEG_1_ database). The results are shown in supplementary Figure S25. By doing this, the meta-band detection only slightly changed, with the blue meta-band still clearly detected around the canonical alpha band. In the second test, the influence of the number of sensors was evaluated. To this end, we constructed two new data subsets using the MEG recordings, matching in the number of sensors and sample size of the EEG_1_ and EEG_2_ databases. As a result, we obtained two new MEG subsets: i) one with 32 sensors and 27 subjects, and ii) other with 19 sensors and 45 subjects. The results are displayed in supplementary Figure S26. It can be appreciated that the detection of the three meta-bands is maintained, although there are some differences that can be due to two factors: i) the reduced sample size (as previously described); and ii) the reduced number of sensors. The influence of the sampling frequency was already assessed as a potential confounding factor in the CMB algorithm. It was observed that for the M/EEG-like synthetic signals, this parameter moderately affected the meta-band detection. This was also analysed with real signals by downsampling the MEG database to 200 Hz (as in the EEG_2_database) and estimating the meta-bands again. The results are shown in supplementary Figure S27 and indicate that the meta-band detection has changed, though the blue meta-band is still clearly detected in the frequencies around the canonical alpha band. However, its presence is less evident than for 1000 Hz. Thus, our findings suggest that the sampling frequency could be, at least partially, influencing the results of the EEG databases. To sum up, these results indicate that these factors (i.e., spatial resolution and sampling frequency) are not sufficient to explain the differences observed between EEG and MEG datasets. In line with this, it can be appreciated that the meta-band detection in the EEG_1_ database is similar to that obtained in the EEG_2_ database, despite the remarkable differences in their spatial resolution (32 *vs.* 19 channels) and sampling frequency (500 *vs.* 200 Hz). Thus, we can conclude that the discrepancies between EEG and MEG are due to the intrinsic dissimilarities between both techniques [2, 4, 103, 6], suggesting an increased sensitivity of MEG to reflect the underlying frequency-dependent network structure.

### 5.4. Limitations and future lines

Although this study yielded interesting and promising results, introducing a new methodology to carry out analyses of the frequential structure of functional brain connectivity, it has some limitations that deserve further discussion.

First of all, we observed a divergence in the results between the MEG and EEG databases. Although several possible reasons to explain the differences have already been discussed, further research is required to find whether it is possible to design a methodology capable of detecting similar meta-bands for EEG signals as those in the MEG database. In line with that, it would also be interesting to evaluate other EEG datasets (with different number of channels and sampling frequencies) using the methodology proposed in this paper. Additionally, different source localization algorithms could be employed to find the extent to which the source inversion method influences the band segmentation achieved by the proposed methodology.

Furthermore, although the introduced methodology is based on a datadriven, automatic algorithm, there are some parameters that could be optimized to adapt specific user requirements (*i.e.*, frequency resolution of the filters, filter order, or sampling frequency, among others). It is likely that the most important parameter is the filter order, with higher values of this parameter providing frequency-accurate solutions (but with an increased presence of noise) and lower values providing smoother solutions.

Moreover, future work should upgrade the implementation of the CMB algorithm to address some of its limitations. In this regard, an interesting new feature would be allowing to configure whether the CMB algorithm accepts meta-bands covering non-adjacent frequency ranges or not. Of note, the implementation of this feature would require prior efforts addressing the issue of spurious meta-band changes (*i.e.*, changes that occur only for isolated frequency bins). Also, the application of the CMB algorithm is now based on the orthogonalized AEC. It has been observed that different connectivity metrics may reflect different neural mechanisms [49, 50]; thereby, in the future, the CMB algorithm could also include new connectivity metrics such as robust phase-based metrics. Additionally, we have based the meta-band segmentation only in the FC patterns. Although these dynamics also reflect, to some extent, information about the oscillatory amplitude, the implementation of the CMB algorithm should be extended to support other input data sources to perform the community detection [25, 26, 104], as power topologies or hybrid data merging FC and power topology information. Furthermore, the CMB algorithm was developed considering the functional connectivity stable across time. This was done due to the high computational cost and complexity associated with considering its dynamic behaviour (dFC approach); nonetheless, in future studies, the CMB algorithm could be updated to support also dFC analyses. Besides, it would also be interesting to merge our methodology with the one previously proposed by Núñez and colleagues, combining time meta-states and frequency meta-bands to provide the time-frequency “building-blocks” of neural dynamics [11, 52].

Finally, the study was conducted with resting-state signals. Further task-related studies should be carried out to assess the potential of the introduced methodology to analyse the changes in the frequential structure of functional brain networks during an structured task.

## 6. Conclusions

In this study, we propose a novel frequency band segmentation approach based on the topology of the functional connectivity matrices in narrow frequency ranges. The data-driven, automatic frequency band segmentation proposed here supports the decomposition into canonical frequency bands at the group level when considering MEG signals; nonetheless, our method is also able to extract the individual idiosyncrasies of frequential structure of connectivity at subject level. Interestingly, we have also observed that the sensitivity of EEG signals to frequency-dependent connectivity neural pattern in to sufficient to identify the underlying meta-band structure.

This study opens the way for personalized, data-driven connectivity analyses, allowing an objective definition of frequency ranges of interest based on the topological similarity of the functional brain network across frequencies. Besides, these analyses, able to quantify the individual idiosyncrasies of neural activity, could be extended to clinical settings to provide a more accurate identification of brain connectivity alterations in several disorders, such as dementia due to Alzheimer’s disease, schizophrenia, depression, or migraine [105, 106, 107, 108]. Furthermore, the CMB algorithm provides a new framework for the characterization of different neural pathologies, as well as to facilitate their diagnosis, thanks to the possibility of studying how they affect to the frequential structure of the functional brain network. The rationale behind the CMB algorithm is in line with the current evolution of science and medicine, which is focused in providing personalized studies, diagnostic tools, and treatments capable of adapting to the particularities of each individual [109, 110].

## Acknowledgments

This research has been funded by “CIBER en Bioingeniería, Biomateriales y Nanomedicina (CIBER-BBN)” through “Instituto de Salud Carlos III” cofunded with ERDF funds. V Rodríguez-González was in receipt of a PIF-UVa grant from the “University of Valladolid”. P. Núñez was funded by the ERA-Net FLAG-ERA JTC2021 project ModelDXConsciousness (Human Brain Project Partnering Project).

