## Supplementary Material for "Connectivity-based Meta-Bands: A new approach for automatic frequency band identification in connectivity analyses"

### 1. GENERATION OF THE SYNTHETIC SIGNALS

#### 1.1. Amplitude-coupled synthetic signals

The pipeline for creating these signals is visually summarized in Figure S1. It is described as follows:

1. Parameters selection:
    a. **Signal length (time)**: 180 seconds.
    b. **Number of channels**: 20 channels.
    c. **Sampling frequency**: 200, 500, 750, and 1000 Hz were tested.
    d. **Noise power**: -19 dBW.
    e. **Carrier amplitude** 1 V.
    f. **Carrier frequency**: Different values were tested for the different tests (from 20 to 40 Hz).
    g. **Modulation index**: 100 %.
    h. **Number of cosines in the modulating signal (*N*)**: A sufficiently high number was selected, to make sure the modulation had a "block" shape in the frequency domain. It was set to 5 times the bandwidth of the modulation (*e.g.*, if the modulation bandwidth is 10 Hz, we will used 50 cosines).
    i. **Modulation bandwidth**: Different values were tested for the different tests (ranging from 0.25 to 40 Hz).
2. Generation of *N* equidistant cosines at different frequencies according to 1.h. In Figure S1 these signals are: $m_1(t), m_2(t) \dots m_N(t)$.
3. Sum all the cosines, thus generating the signal $m(t)$.
4. Generate the AM modulation using the signal generated in step 3 as the modulating signal. The resulting signal is called $s(t)$.
5. Compute the product of the AM-modulated signals with specific weights to generate a coupling pattern. These weights are called $w_1, w_2 \dots w_N$.
6. Add white gaussian noise ($o_1, o_2 \dots o_N$ in Figure S1) to the signals in step 5. The resulting signals for each channel are named $s_1(t), s_2(t) \dots s_N(t)$.

If the signal is composed by more than one AM modulation, the steps 4-6 will be repeated for each of the AM modulations composing the AM signal, and the resulting signals will be added up. One example of the power spectral density and time course of an amplitude-coupled synthetic signal with two AM modulations is depicted in Figure S2.

To separate the different AM modulations in one signal, clearly-defined coupling patterns between channels were used by multiplying the signal, $s(t)$, by the binary weights ($w_1, w_2 \dots w_N$). Specifically, we generated the following channel patterns: i) channels 1 to 5 multiplied by 1, remaining channels multiplied by 0; and ii) channels 15 to 20 multiplied by 1, remaining channels multiplied by 0. In most of the scenarios, the first pattern is used. The second pattern is used only when the signal is composed of two AM modulations. These channel patterns result in specific connectivity matrices that can be observed in Figure S3.

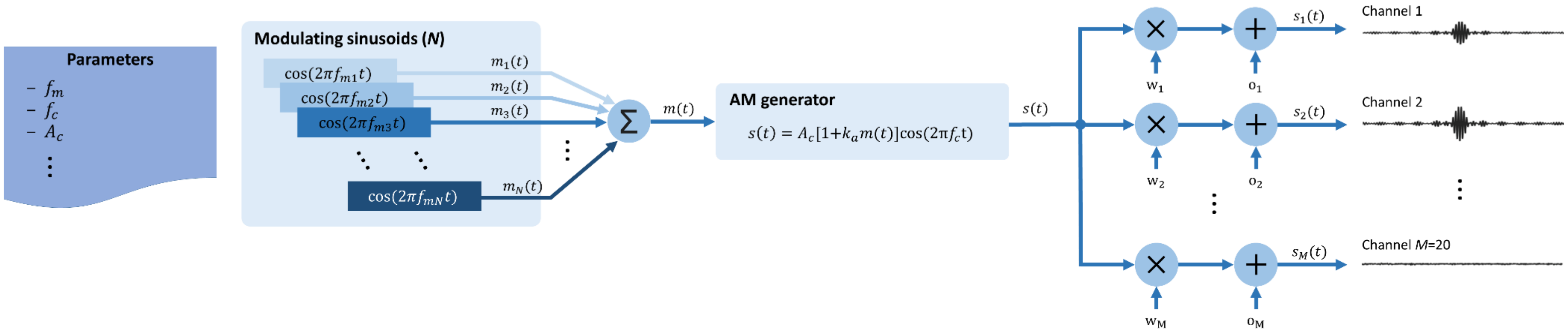

***Figure S1***. *Graphical pipeline for the generation of the amplitude-coupled synthetic signals*

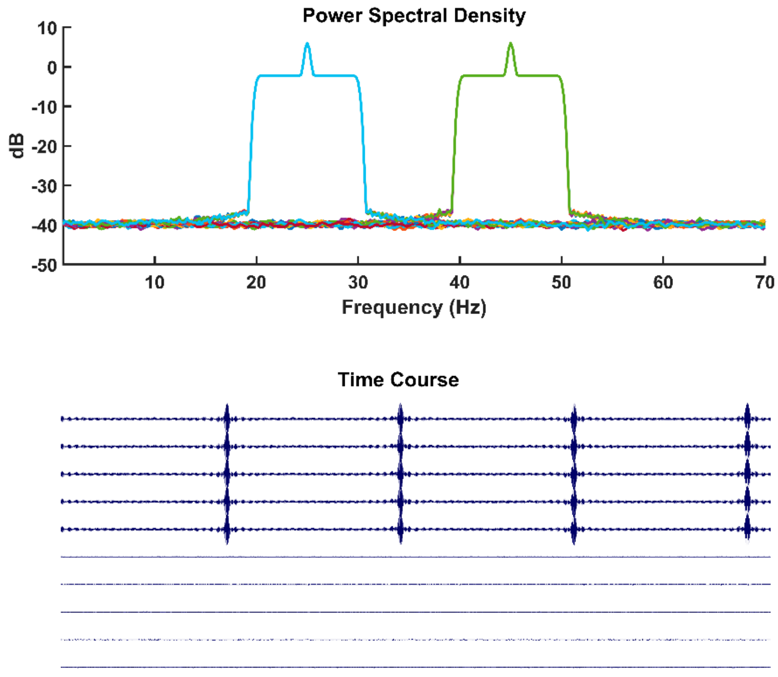

***Figure S2***. *Example of power spectral density and time course of an amplitude-coupled synthetic signal.*

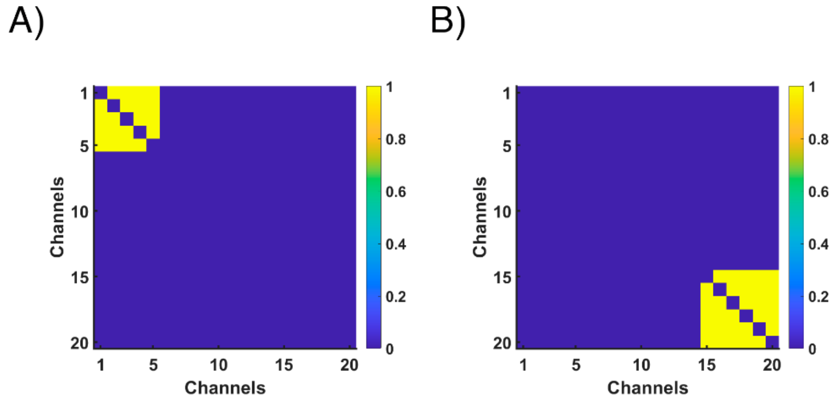

***Figure S3***. *Connectivity matrices for the amplitude-coupled synthetic signals*

#### 1.2. M/EEG-like synthetic Signals

The pipeline for creating these signals is visually summarized in Figure S4. It is described as follows:

1. Parameters selection:
    a. **Number of frequency bands**: From 1 to 6.
    b. **Frequency range of the bands**: Different values were tested (see table S1).
    c. **Overlap**: Both "*Continuous*" and "*Not Continuous*" frequency overlap patterns were tested. In this scenario "*Continuous*" means not leaving any gap between meta-bands.
2. Calculate frequency-dependent connectivity matrices at source level (68 ROIs) based on the orthogonalized AEC for 100 MEG recordings: $\mathbf{A}_1, \mathbf{A}_2 \dots \mathbf{A}_{100}$.
3. For each of these frequency-dependent connectivity matrices (excluding the diagonal values), estimate the probability density function (pdf) of a "real" coupling values.
4. Build a surrogate signal from a real MEG recording by means of the AAFT method, obtaining the signal $\mathbf{X}(t)$.
5. Filter the signal in the frequency range of interest. This filtered signal is called $\mathbf{X}_1^{\mathrm{b}}(t)$.
6. Use the pdf from step 3 to generate random values that follow the distribution of the "real" coupling values by applying the inverse transform sampling methodology. Distribute these values randomly into a connectivity matrix, thus obtaining a pseudo-real (with a "real" coupling distribution) connectivity matrix. This connectivity matrix is called $\mathbf{A}_1^{\mathrm{S}}$
7. Generate the coupled signal using the AAFT signal (signal from step 4) and the connectivity matrix (step 6). Each source will be the sum of all the sources weighted using the adjacency matrix. Z-score this signal to smooth the transitions between bands and noise in the final signal. This band-limited coupled signal is named $\mathbf{X}_1^{\mathrm{c}}(t)$.
8. Repeat steps 5-7 for all the frequency bands to define. Thus, we obtained the signals: $\mathbf{X}_1^{\mathrm{c}}(t), \mathbf{X}_2^{\mathrm{c}}(t), \dots \mathbf{X}_{\mathrm{N}}^{\mathrm{c}}(t)$, with *N* the number of bands.
9. Band-stop filter the signal resulting from the AAFT (signal from step 4) in the bandwidth of all the meta-bands defined. Z-score this signal to smooth the transitions between states and noise in the final signal. This band-stop filtered signal is referred as $\mathbf{X}_{\mathrm{u}}(t)$
10. Sum the signal obtained in step 9 with all the signals generated in the different iterations of step 8. The final signal will be: $\mathbf{S}(t) = \mathbf{X}_{\mathrm{u}}(t) + \mathbf{X}_1^{\mathrm{c}}(t) + \mathbf{X}_2^{\mathrm{c}}(t) + \cdots + \mathbf{X}_{\mathrm{N}}^{\mathrm{c}}(t)$

In this case, the synthetic frequency-dependent connectivity matrices are more similar to real neurophysiological signals than the synthetic signals generated in the previous section. An example of the power spectral density and time course of one of these signals is displayed in Figure S5. Additionally, an example of a synthetic connectivity matrix resulting from this method can be observed in Figure S6.

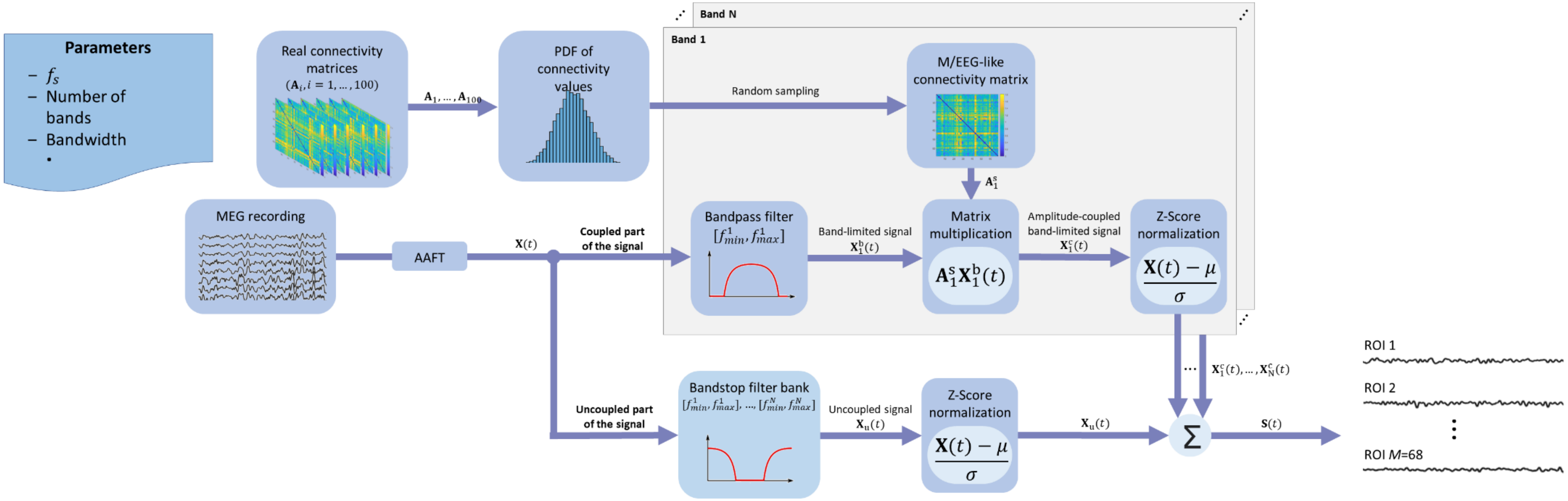

***Figure S4***. *Graphical pipeline for the generation of the M/EEG-like synthetic signals*

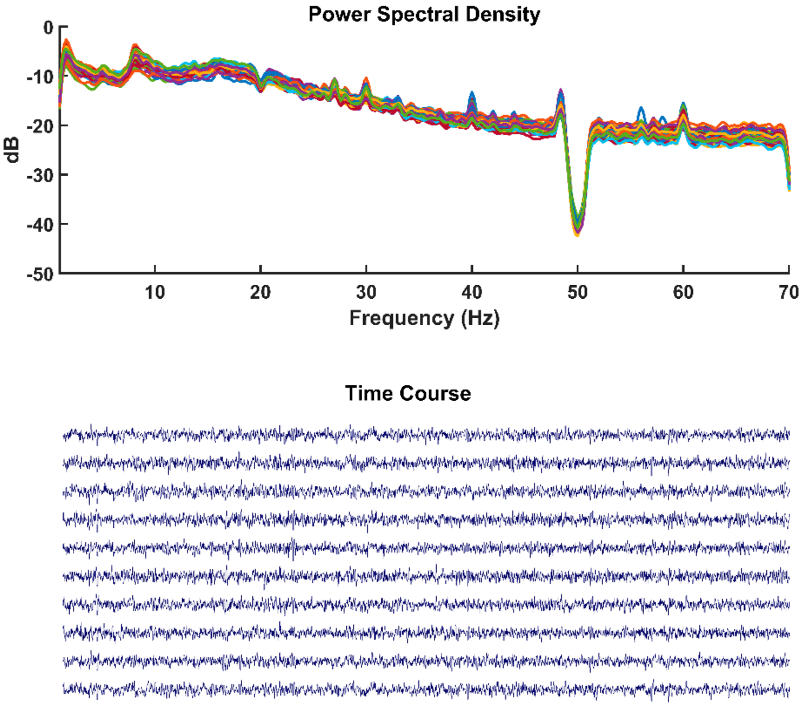

***Figure S5***. *Example of power spectral density and time course of a M/EEG-like synthetic signal.*

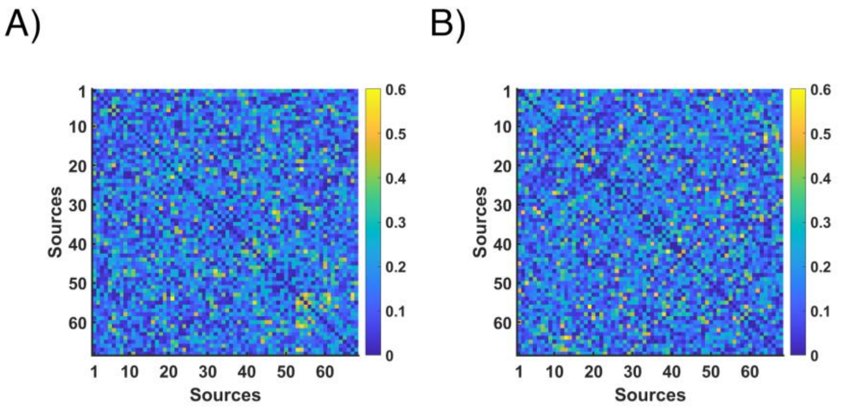

***Figure S6***. *A) Example of connectivity matrix from a real MEG signal. B) Connectivity matrix from a M/EEG-like synthetic signals*

***Table S1***. *Meta-bands defined for the M/EEG-like synthetic signals*

| Number of Meta-Bands | Filled (Hz) | Not filled (Hz) |
|---|---|---|
| *1* | 1-70 | 5-20 |
| *2* | 1-20, 20-70 | 5-15, 30-45 |
| *3* | 1-20, 20-50, 50-70 | 3-13, 25-35, 55-68 |
| *4* | 1-15, 15-30, 30-45, 45-70 | 3-12, 18-25, 30-45, 55-68 |
| *5* | 1-10, 10-25, 25-35, 35-45, 45-70 | 3-10, 15-22, 25-32, 38-45, 55-68 |
| *6* | 1-10, 10-25, 25-35, 35-45, 45-60, 60-70 | 3-10, 15-22, 25-32, 38-45, 52-58, 60-68 |

### 2. RESULTS FOR THE SYNTHETIC SIGNALS

#### 2.1. Assessing the CMB algorithm with amplitude-coupled synthetic signals

The amplitude-coupled synthetic signals provide a framework to perform extensive tests on the CMB algorithm, thus allowing to assess multiple parameters in a computational cost-effective way. In the following sections, the results from analyzing the influence of different parameters in the results achieved by the CMB algorithm are described. For these analyses, all the epochs of each signal were stacked to perform the community detection. Due to the wide-sense stationarity property of these signals, averaging across trials was not required.

The parameters were assessed by varying one while keeping the others to their default values. The default values were: sampling frequency of 1000 Hz, a single underlying AM of 10 Hz bandwidth (from 30 to 40 Hz), filter order of 500, and frequency resolution of 1 Hz.

##### 2.1.1. Influence of the filter order in the meta-bands detected

The filter order is one of the parameters that most influences the CMB algorithm. The analyzed filter order values were: 100, 250, 500, 750, 1000, 1250, 1500, 2000, 2500, and 3000. The analyses were performed with a sampling frequency of 1000 Hz and a single AM with a bandwidth of 10 Hz (from 30 to 40 Hz). The results for these analyses are depicted in Figure S7. There, it can be appreciated that for all the filter orders, the recovered connectivity matrix is similar to the one originally generated. On the one hand, lower filter orders yield a decreased frequency resolution to detect the underlying meta-band (*i.e.*, lower efficiency detecting the edges of the meta-band). On the other hand, higher filter orders lead to a decreased performance of the algorithm, with a noisier detection in the frequencies where no meta-band was specified and, thus, less defined connectivity matrices with lower coupling values. Besides, higher filter orders have an increased computational burden. In consequence, we have employed a filter order of 500, as it provides a good balance between the aforementioned effects for our purpose.

It is difficult to provide specific rules for determining that balance, as there are many influencing factors that totally depend on the specific idiosyncrasies of the study. Some applications may require maximizing the spectral resolution, even with the penalty of noisier detection (that could be smoothed afterwards); in that case, a high filter order may be selected (filter order ≥ 1500). Other applications may prefer to have smoother results, as the specific limits of the meta-bands are not considered so important, thus selecting a lower filter order (filter order ≤ 500). Also, in other studies the computational resources available may also be a limitation to be considered, with lower values of filter order involving less computational cost. A good rule of thumb could be to evaluate different filter orders on a subset of the original sample, and based on that, decide which one best suits the specific need of the study. Based on our experience, a good starting point for tuning the filter order would be between 500 and 1000.

It has to be mentioned that, as depicted in **Error! Reference source not found.**.A, for a filter order of 100, the underlying frequency band was properly recovered, but split in two different meta-bands with a very similar network topology (and similar to the original one). This phenomenon can also be observed in other scenarios (see Figure S8.F, Figure S8.G ). This issue only emerges when the following two conditions are met: i) there is only one meta-band, ii) that band is wider than approximately 30 Hz. Results in Figure S12, Figure S14, Figure S16, and Figure S18 (in the “2.2. Assessing the CMB algorithm with M/EEG-like synthetic signals” section) support this finding, as the band-splitting phenomenon occurs when there is only one underlying meta-band (panels A in the Figures), but not when there are at least two, one of them being 50 Hz-wide (panels B-F in the Figures). Although this issue is a mathematical restriction of the methodology, it only occurs in a specific scenario very unlikely to happen in real-life, so it has not been included as a limitation in the "5.4. Limitations and Future Lines" section.

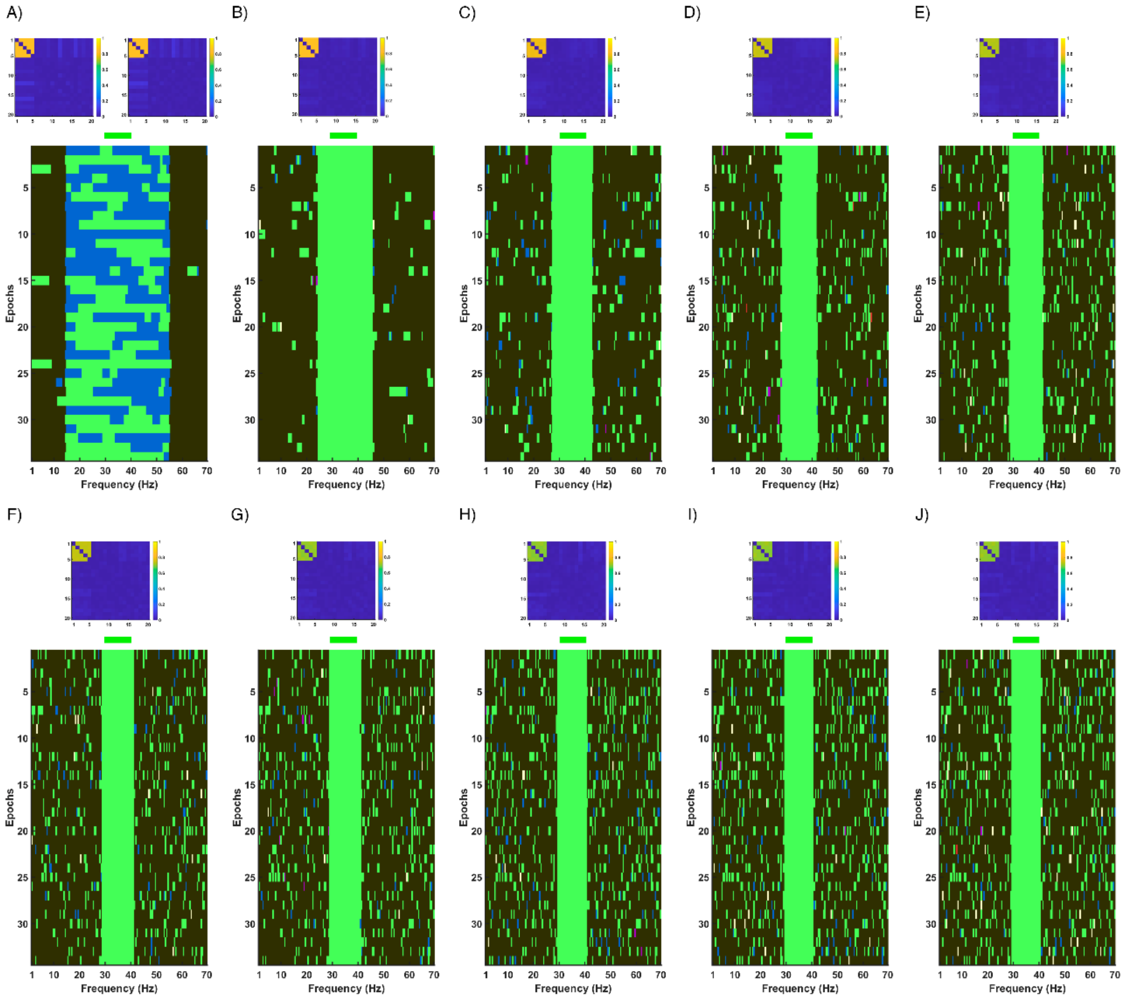

*Figure S7. Meta-bands detected for synthetic signals (sampling frequency of 1000 Hz) for different filter orders: A) 100, B) 250, C) 500, D) 750, E) 1000, F) 1250, G) 1500, H) 2000, I) 2500, and J) 3000. The upper plot is the connectivity matrix of the modal meta-band detected in the frequencies where the original meta-band was defined. If two meta-bands are clearly dominating in these frequencies, the connectivity matrices corresponding with both meta-bands are depicted. Finally, in the lower plot, the Y-axis represents each epoch from the original signal, while X-axis depicts each frequency bin. Different colors represent different communities. Of note, the same color may be associated to different communities across figures. Above these plots, there is a green bar indicating the frequencies where the original meta-band was defined.*

##### 2.1.2. Influence of bandwidth in the meta-bands detected

The meta-bands detected for different bandwidths of the underlying meta-bands were tested. The aim was to assess whether there exists a lower or upper bound in the bandwidth of the meta-bands that the CMB algorithm is able to detect. In Figure S8, the meta-bands detected for underlying meta-bands with bandwidth of 1, 2, 5, 10, 20, 30 and 40 Hz can be observed.

It can be appreciated that the ability of the algorithm to detect narrow meta-bands is influenced by two factors. The first one is the filter order, because as we have seen in previous section, lower filter orders limit the frequency resolution of the meta-band detection. The other factor is the frequency resolution of the filters (*i.e.*, their bandwidth) selected to generate the meta-bands. With a high enough filter order, the limit is the frequency resolution of the filters, as the CMB algorithm is capable of properly recovering 1 Hz meta-bands (when constructed with a frequency resolution of the filters of 1 Hz), as it can be appreciated in **Error! Reference source**

**not found.**. Deeper insights on the frequency resolution of the filters will be described in following sections.

Moreover, wider bandwidths provide better-defined connectivity matrices, as there are more frequency bins with that specific connectivity pattern. Furthermore, starting from a bandwidth of 30 Hz, the underlying meta-band is segmented into different meta-bands (as happened in the previous section in the scenario of a filter order of 100).

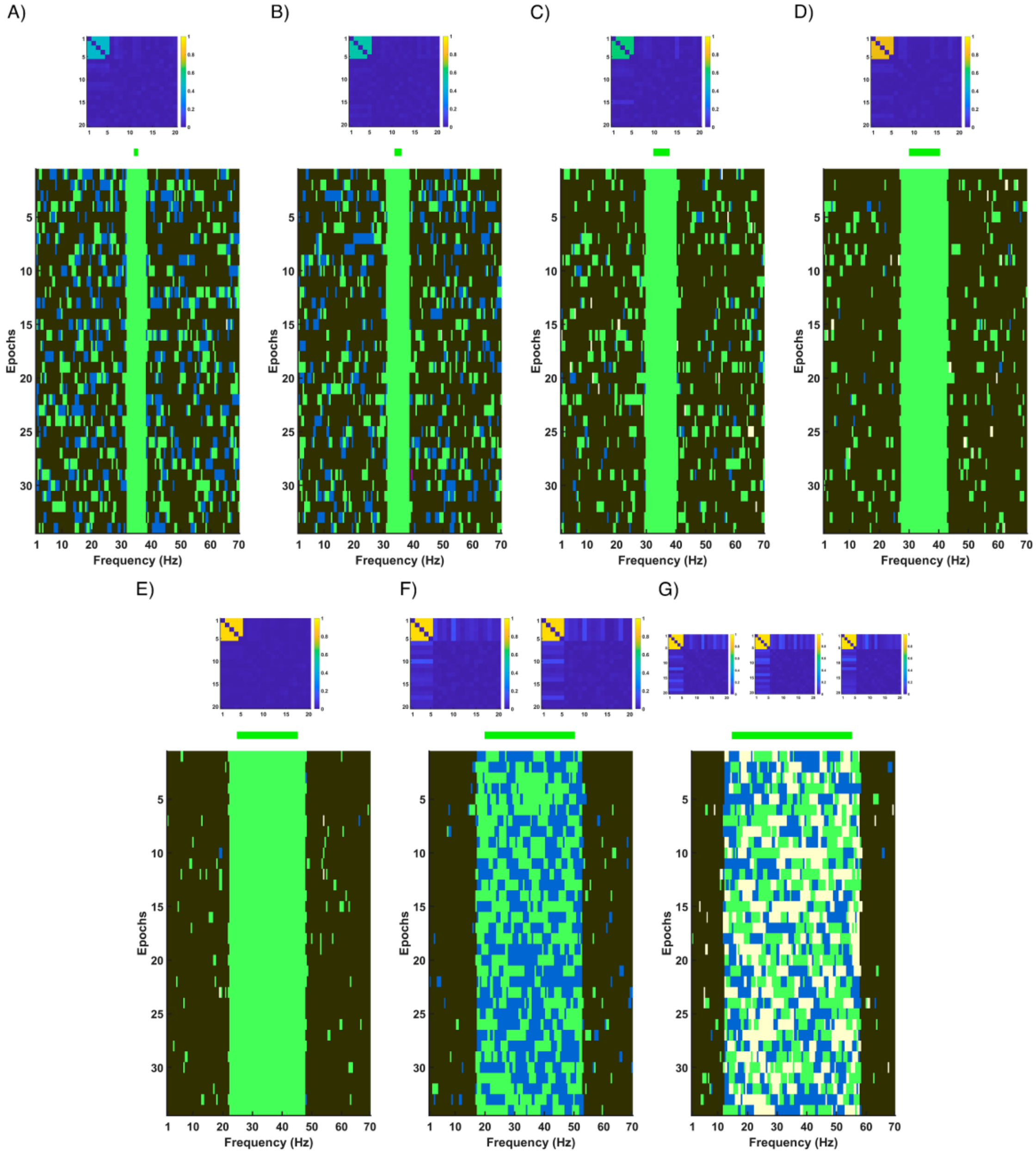

*Figure S8. Meta-bands detected for synthetic signals (sampling frequency of 1000 Hz) for different bandwidths of the underlying meta-band: A) 1 Hz, B) 2 Hz, C) 5 Hz, D) 10 Hz, E) 20 Hz, F) 30 Hz, and G) 40 Hz. The upper plot is the connectivity matrix of the modal meta-band detected in the frequencies where the original meta-band was defined. If more than one meta-band are clearly dominating in these frequencies, the connectivity matrices corresponding with all these meta-bands are depicted. Finally, in the lower plot, the Y-axis represents each epoch from the original signal, while X-axis depicts each frequency bin. Different colors mean different communities. Of note, the same color may be associated to different communities across figures. Above these plots, there is a green bar indicating the frequencies where the original meta-band was defined.*

##### 2.1.3. Influence of sampling frequency in the meta-bands detected

Meta-bands were also calculated using synthetic signals with different sampling frequencies, obtained by downsampling the original 1000 Hz signals. The signals used in this section have a single underlying meta-band of 10 Hz bandwidth (from 30 to 40 Hz). In Figure S9, the meta-bands detected for different sampling frequencies (200, 500, 750, and 1000 Hz) are depicted. There, it can be observed that lower sampling frequencies provide slightly less widening in the bands for the same filter order. Besides the lower the sampling frequency, the lower regularity in the detection (*i.e*, spurious state changes) in frequencies where no underlying band was defined.

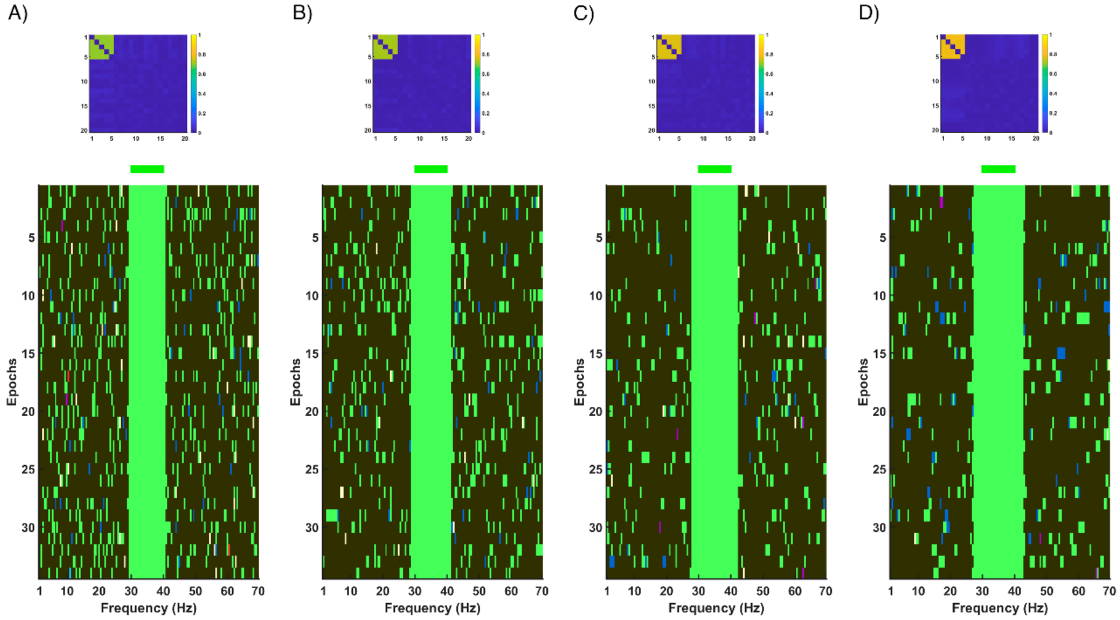

*Figure S9. Meta-bands detected for synthetic signals (filter order of 500) for different sampling frequencies: A) 200 Hz, B) 500 Hz, C) 750 Hz, and D) 1000 Hz. The upper plot is the connectivity matrix of the modal meta-band detected in the frequencies where the original meta-band was defined. Finally, in the lower plot, the Y-axis represents each epoch from the original signal, while X-axis depicts each frequency bin. Different colors mean different communities. Of note, the same color may be associated to different communities across figures. Above these plots, there is a green bar indicating the frequencies where the original meta-band was defined.*

##### 2.1.4. Ability of the CMB algorithm to detect two overlapping bands

Other relevant feature of the CMB algorithm to assess is whether it is able to detect contiguous (or even overlapping) underlying meta-bands. In this regard, three different scenarios are depicted in Figure S10 (from left to right): i) two meta-bands separated by 6 Hz; ii) two contiguous meta-bands; and iii) two partially overlapping meta-bands. It can be appreciated that the CMB algorithm is capable of detecting close, contiguous, and even overlapping meta-bands. In this latter scenario, in the frequency ranges where two meta-bands are overlapped, only one of them is detected.

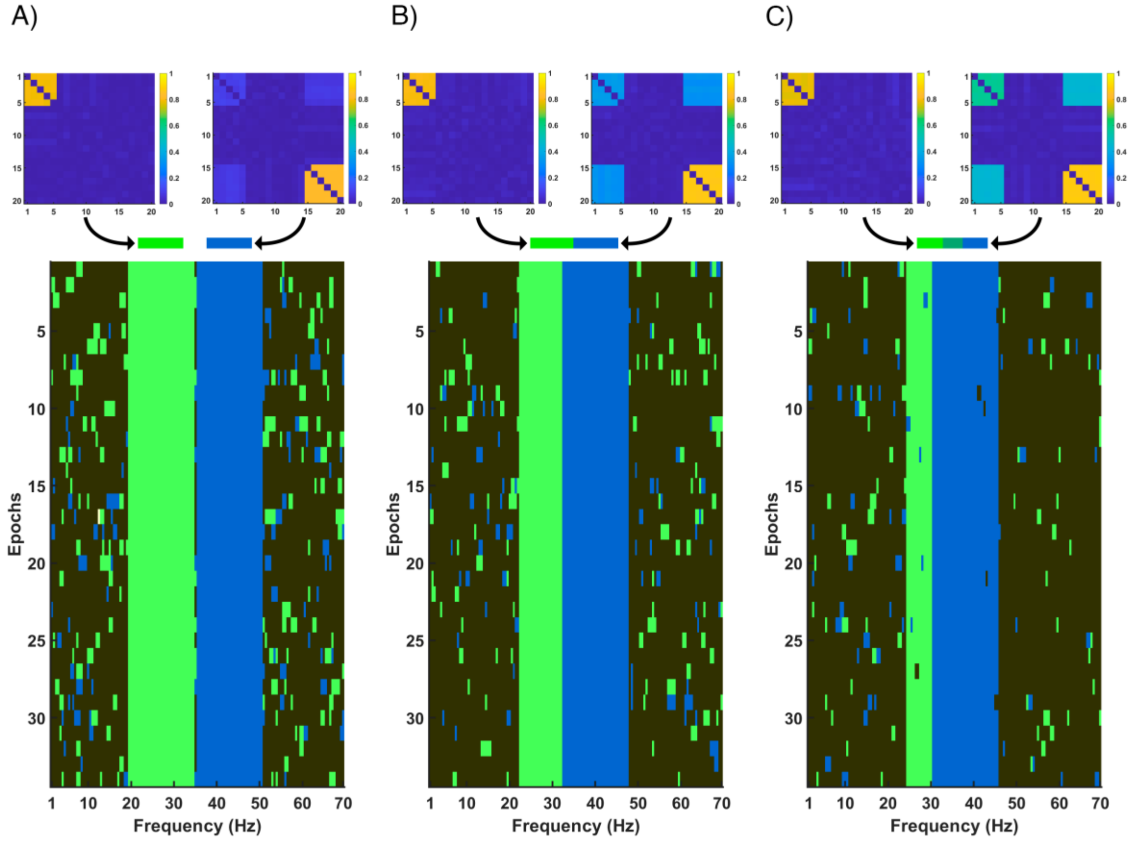

*Figure S10. Meta-bands detected for synthetic signals (sampling frequency of 1000 Hz) when two underlying meta-bands are present: A) separated by 6 Hz, B) contiguous, and C) overlapping. The upper plot displays the connectivity matrices of the modal meta-bands detected in the frequencies where the original meta-bands were defined. In the lower plot, the Y-axis represents each epoch from the original signal, while the X-axis depicts each frequency bin. Different colors represent different communities detected by the algorithm. Of note, the same color can be associated to unrelated communities across figures. Above these plots, the green and blue bars indicate the frequencies where the original meta-bands were defined.*

##### 2.1.5. Influence of frequency resolution in the meta-bands detected

In order to assess the lower limit of the CMB algorithm to detect narrow underlying meta-bands, frequency resolutions of 0.5 and 0.25 Hz were also tested. It is noteworthy that increasing the frequency resolution significantly increases the computational burden of the CMB algorithm, thus it has to be considered whether it is worth doing so for each specific user application. Due to the high computational cost, the number of iterations used in the execution of the Louvain GJA method was decreased from 250 to 100, which has been proven to still be enough for the algorithm to converge (Núñez et al., 2021).

The meta-bands detected for different combinations of frequency resolution (0.5 and 0.25 Hz) and filter order (1500, 2000, 2500, and 3000) are depicted in Figure S11. It can be observed that for a frequency resolution of 0.5 Hz (panels A-D), the CMB algorithm is able to detect meta-bands with a bandwidth of 0.5 Hz (although high-order filtering is required). In addition, it can also be appreciated that with a frequency resolution of 0.25 Hz (panels E-H), the CMB algorithm is not able to recover meta-bands with a bandwidth of 0.25 Hz.

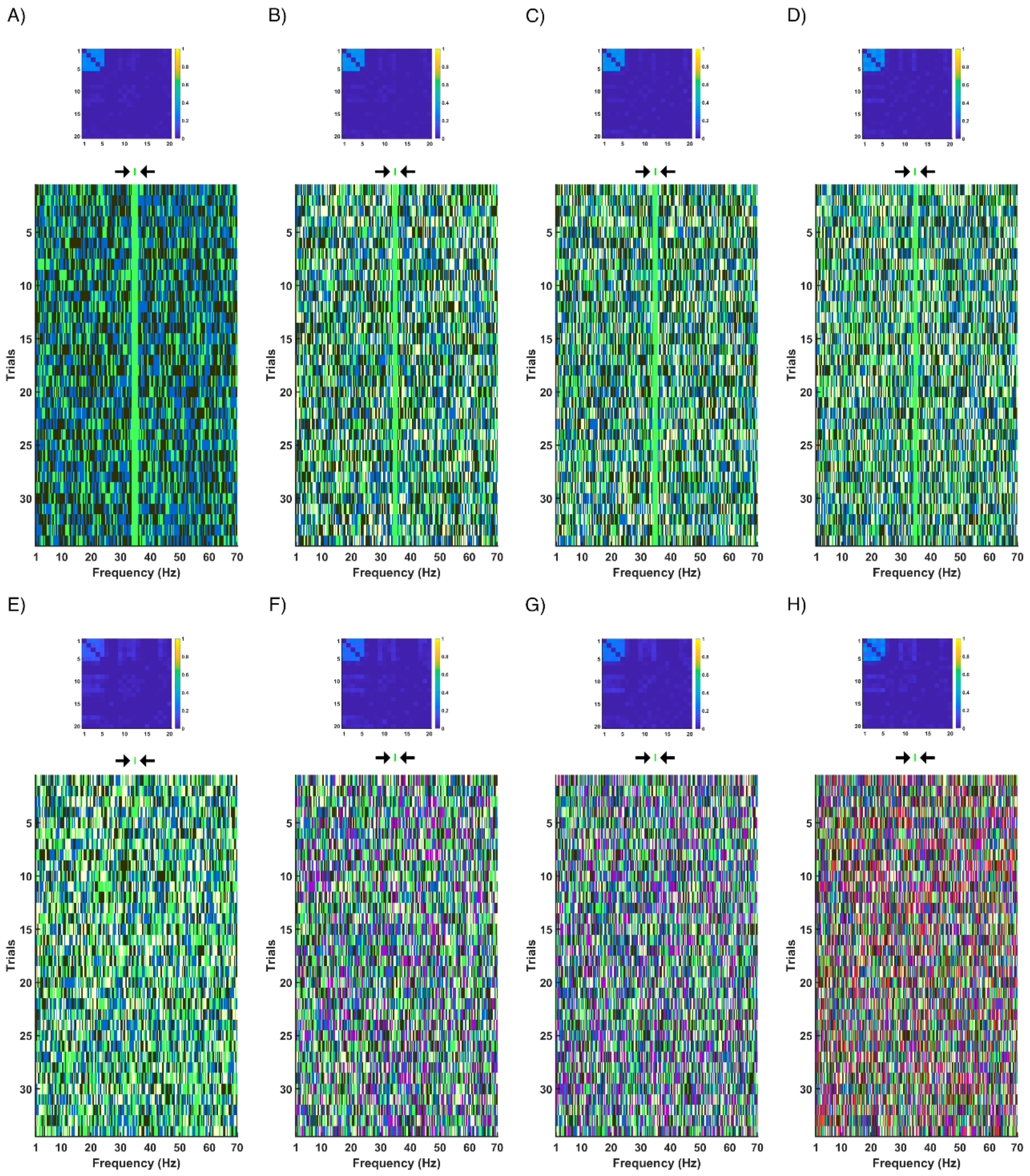

*Figure S11. Meta-bands detected for synthetic signals (sampling frequency of 1000 Hz) using low frequency resolutions: A-D) 0.5 Hz bandwidth with a frequency resolution of 0.5 Hz, E-H) 0.25 Hz bandwidth with a frequency resolution of 0.25 Hz; and different filter orders: A) and E) 1500, B) and F) 2000, C) and G) 2500, and D) and H) 3000. The upper plot displays the connectivity matrix of the modal meta-band detected in the frequencies where the original meta-band was defined. Finally, in the lower plot, the Y-axis represents each epoch from the original signal, while the X-axis depicts each frequency bin. Different colors represent different communities. Of note, the same color may be associated to different communities across figures. Above these plots, there is a green bar indicating the frequencies where the original meta-band was defined. For plots E-H) the CMB algorithm was not able to detect the underlying meta-bands.*

#### 2.2. Assessing the CMB algorithm with M/EEG-like synthetic signals

The amplitude-coupled synthetic signals suffer from the limitation of not providing a real-life scenario. Therefore, M/EEG-like synthetic signals with known ground-truth (*i.e.*, the underlying meta-bands) were also generated. With these signals, it was tested whether the CMB algorithm is capable of recovering a variable number of underlying meta-bands. This was tested for two different underlying meta-band configurations: i) "Continuous", letting the bands fill all the spectrum, and ii) "Not continuous", leaving frequency ranges between meta-bands without defining an underlying network topology. Moreover, the influence of the sampling frequency in the CMB algorithm was assessed by repeating these analyses with three additional sampling frequencies (apart from the original 1000 Hz): 200, 500, and 750 Hz.

##### 2.2.1 Ability of the CMB algorithm to detect from 1 to 6 meta-bands in the “Continuous” scenario

Figures S12, S14, S16, and S18 depicts the meta-bands detected when: A) one, B) two, C) three, D) four, E) five, and F) six underlying meta-bands are generated for the “Continuous” scenario (sampling frequency of: Figure S12, 1000 Hz; Figure S14, 750 Hz; Figure S16, 500 Hz; and Figure S18, 250 Hz). It can be observed that the underlying meta-bands are properly recovered, with the topologies of the detected bands being very similar to the ground-truth underlying bands. Of note, when only one meta-band is generated, the algorithm segments the underlying band into three different bands, although the topology is still similar to the generated with only one meta-band.

##### 2.2.2 Ability of the CMB algorithm to detect from 1 to 6 meta-bands in the “Not continuous” scenario

Figures S13, S15, S17, and S19 depicts the meta-bands detected when: A) one, B) two, C) three, D) four, E) five, and F) six underlying meta-bands are generated for the “Not continuous” scenario (sampling frequency of: Figure S13, 1000 Hz; Figure S15, 750 Hz; Figure S17, 500 Hz; and Figure S19, 250 Hz). In this case, all the underlying meta-bands are properly recovered, with the topologies of the detected bands being very similar to the generated meta-bands.

##### 2.2.3 Influence of sampling frequency in the meta-bands detected

In the previous figures (Figures S12-S19), the influence of the sampling frequency in the meta-band detection in this M/EEG-like scenario can be appreciated. On the one hand, the topologies of the detected meta-bands are successfully recovered for all the assessed sampling frequencies. On the other hand, the FAS presents some differences across sampling frequencies, with lower values of sampling frequencies yielding detections with an increased number of state changes in the “uncoupled” frequency ranges (*i.e.*, the frequency ranges where no underlying meta-band was defined).

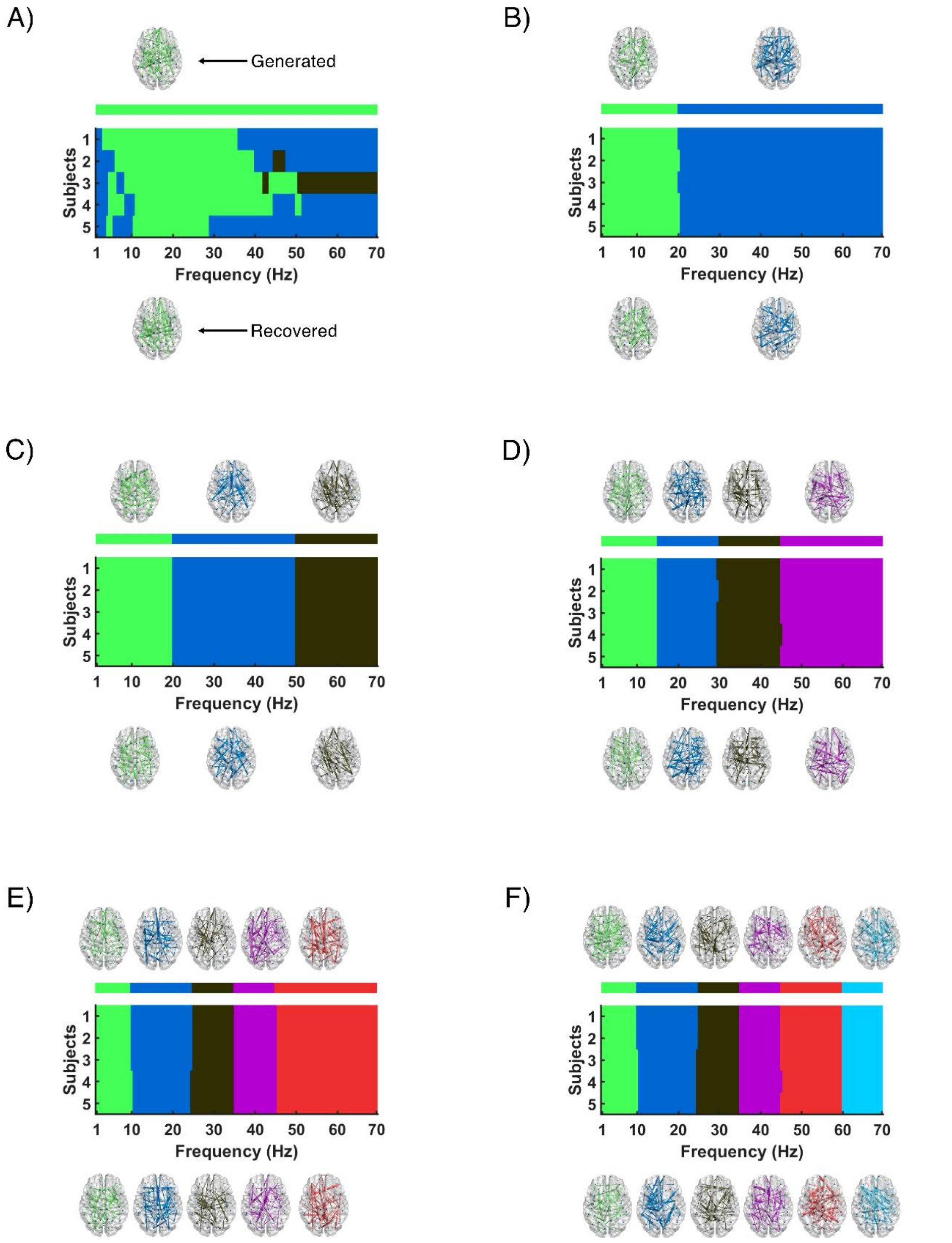

*Figure S12. Meta-bands detected with M/EEG-like synthetic signals for different numbers of underlying bands for the "Continuous" scenario (sampling frequency = 1000 Hz): A) 1, B) 2, C) 3, D) 4, E) 5, and F) 6 meta-bands. The upper brain plots display the network topologies used to generate the underlying meta-bands (i.e., the ground-truth network topologies), while the brain plots on the bottom row depict the network topologies of the detected meta-bands. In the central plot, the Y-axis represents the subjects, while the X-axis depicts each frequency bin. Above this plot, there is a bar indicating the frequencies where each of the original meta-bands were defined. Different colors represent different communities. The original and detected brain topologies and its corresponding meta-band are depicted with the same color. Of note, the same color can be associated to different communities across figures.*

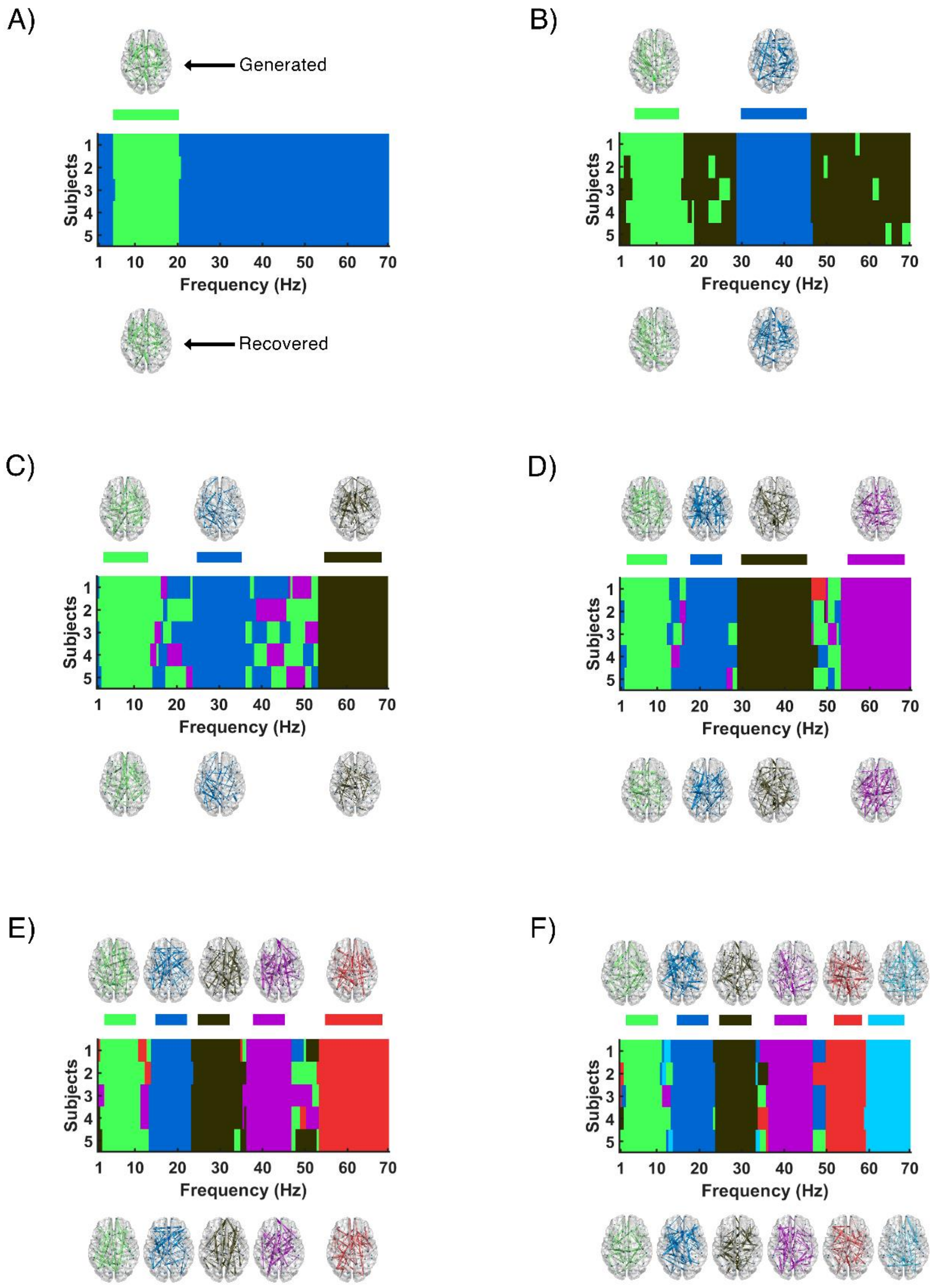

*Figure S13. Meta-bands detected with M/EEG-like synthetic signals for different numbers of underlying bands for the "Not continuous" scenario (sampling frequency = 1000 Hz): A) 1, B) 2, C) 3, D) 4, E) 5, and F) 6 meta-bands. The upper brain plots display the network topologies used to generate the underlying meta-bands (i.e., the ground-truth network topologies), while the brain plots on the bottom row depict the network topologies of the detected meta-bands. In the central plot, the Y-axis represents the subjects, while the X-axis depicts each frequency bin. Above this plot, there is a bar indicating the frequencies where each of the original meta-bands were defined. Different colors represent different communities. The original and detected brain topologies and its corresponding meta-band are depicted with the same color. Of note, the same color can be associated to different communities across figures.*

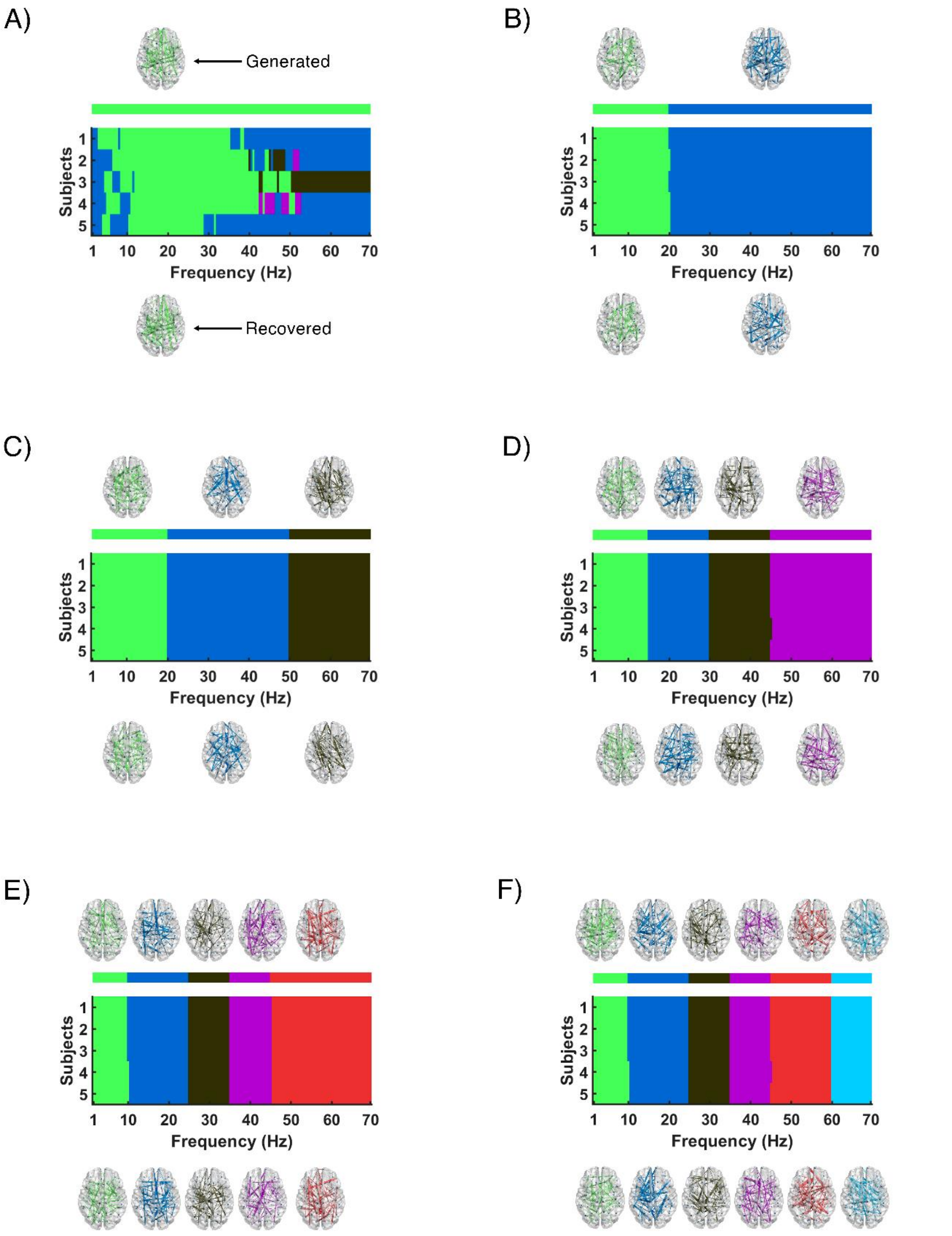

*Figure S14. Meta-bands detected with M/EEG-like synthetic signals for different numbers of underlying bands for the "Continuous" scenario (sampling frequency = 750 Hz): A) 1, B) 2, C) 3, D) 4, E) 5, and F) 6 meta-bands. The upper brain plots display the network topologies used to generate the underlying meta-bands (i.e., the ground-truth network topologies), while the brain plots on the bottom row depict the network topologies of the detected meta-bands. In the central plot, the Y-axis represents the subjects, while the X-axis depicts each frequency bin. Above this plot, there is a bar indicating the frequencies where each of the original meta-bands were defined. Different colors represent different communities. The original and detected brain topologies and its corresponding meta-band are depicted with the same color. Of note, the same color can be associated to different communities across figures.*

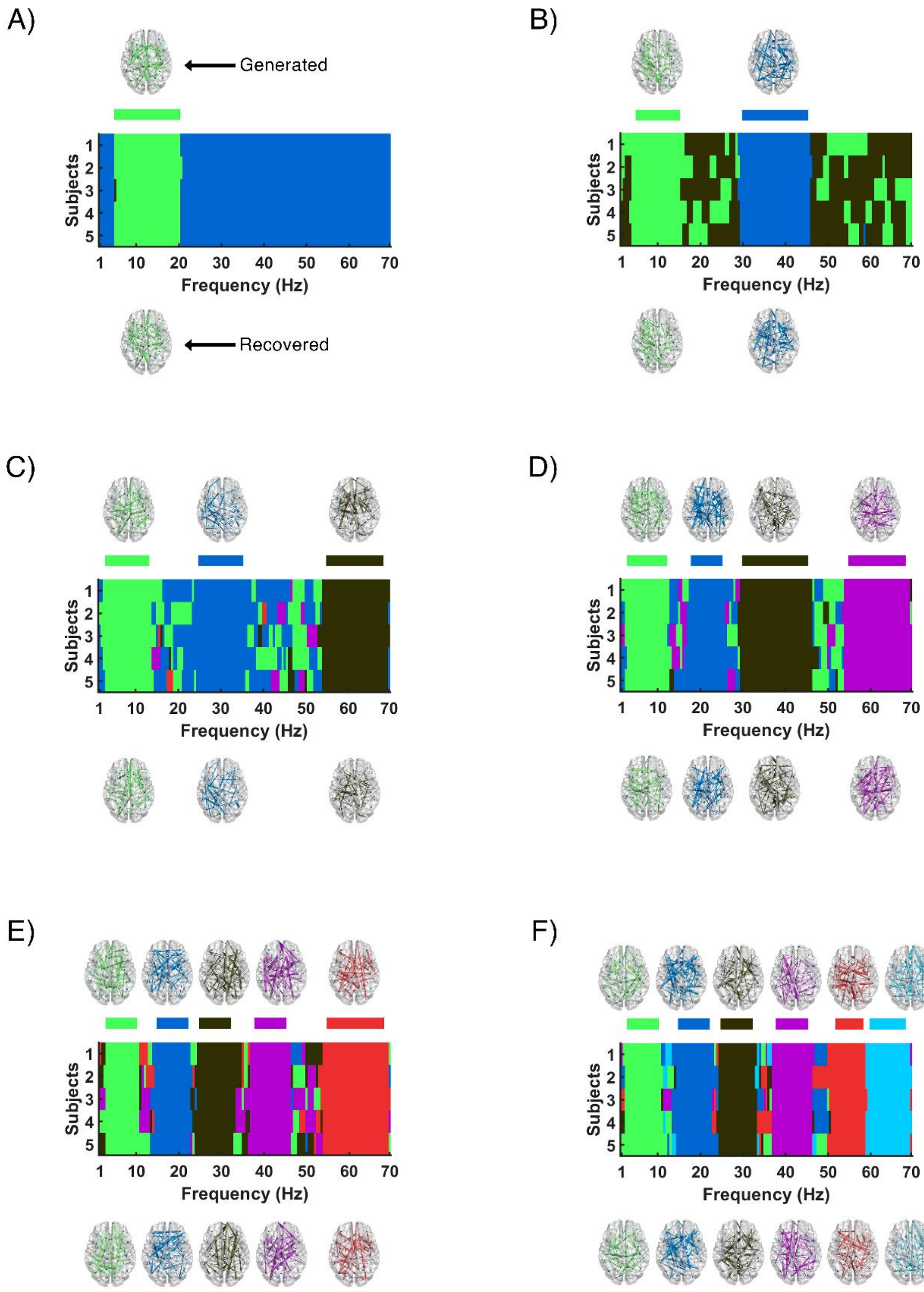

*Figure S15. Meta-bands detected with M/EEG-like synthetic signals for different numbers of underlying bands for the "Not continuous" scenario (sampling frequency = 750 Hz): A) 1, B) 2, C) 3, D) 4, E) 5, and F) 6 meta-bands. The upper brain plots display the network topologies used to generate the underlying meta-bands (i.e., the ground-truth network topologies), while the brain plots on the bottom row depict the network topologies of the detected meta-bands. In the central plot, the Y-axis represents the subjects, while the X-axis depicts each frequency bin. Above this plot, there is a bar indicating the frequencies where each of the original meta-bands were defined. Different colors represent different communities. The original and detected brain topologies and its corresponding meta-band are depicted with the same color. Of note, the same color can be associated to different communities across figures.*

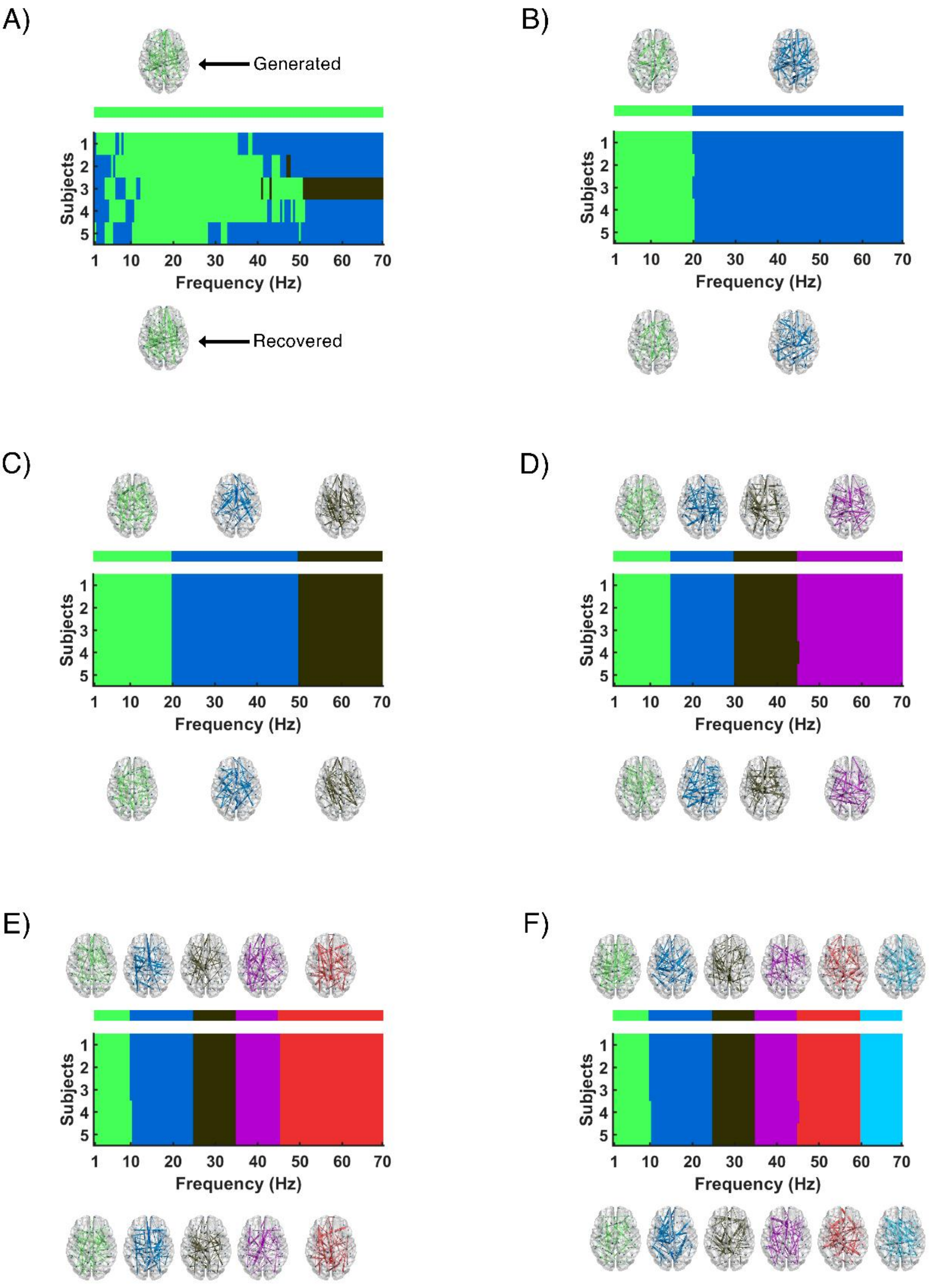

*Figure S16. Meta-bands detected with M/EEG-like synthetic signals for different numbers of underlying bands for the "Continuous" scenario (sampling frequency = 500 Hz): A) 1, B) 2, C) 3, D) 4, E) 5, and F) 6 meta-bands. The upper brain plots display the network topologies used to generate the underlying meta-bands (i.e., the ground-truth network topologies), while the brain plots on the bottom row depict the network topologies of the detected meta-bands. In the central plot, the Y-axis represents the subjects, while the X-axis depicts each frequency bin. Above this plot, there is a bar indicating the frequencies where each of the original meta-bands were defined. Different colors represent different communities. The original and detected brain topologies and its corresponding meta-band are depicted with the same color. Of note, the same color can be associated to different communities across figures.*

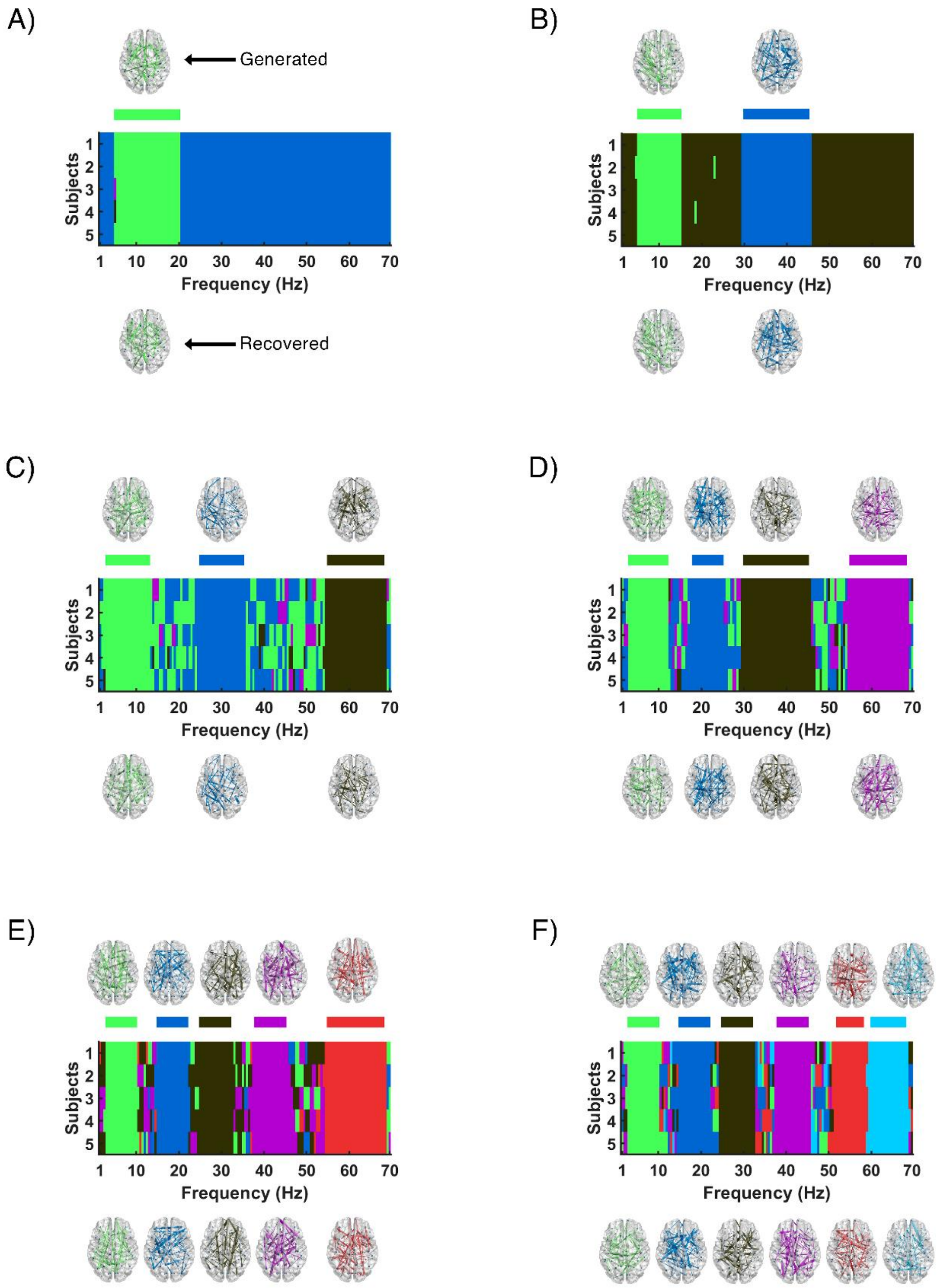

*Figure S17. Meta-bands detected with M/EEG-like synthetic signals for different numbers of underlying bands for the "Not continuous" scenario (sampling frequency = 500 Hz): A) 1, B) 2, C) 3, D) 4, E) 5, and F) 6 meta-bands. The upper brain plots display the network topologies used to generate the underlying meta-bands (i.e., the ground-truth network topologies), while the brain plots on the bottom row depict the network topologies of the detected meta-bands. In the central plot, the Y-axis represents the subjects, while the X-axis depicts each frequency bin. Above this plot, there is a bar indicating the frequencies where each of the original meta-bands were defined. Different colors represent different communities. The original and detected brain topologies and its corresponding meta-band are depicted with the same color. Of note, the same color can be associated to different communities across figures.*

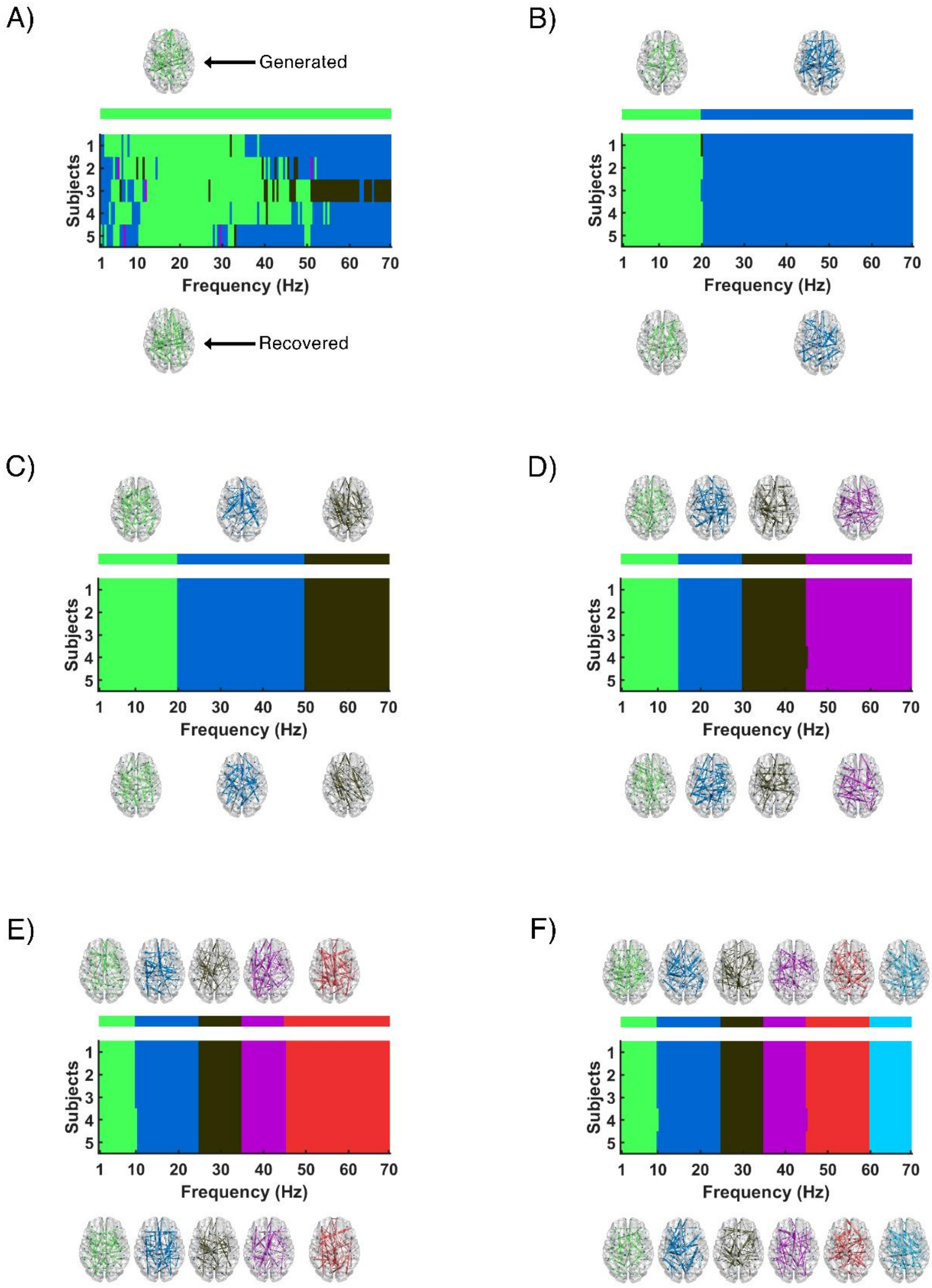

*Figure S18. Meta-bands detected with M/EEG-like synthetic signals for different numbers of underlying bands for the “Continuous" scenario (sampling frequency = 200 Hz): A) 1, B) 2, C) 3, D) 4, E) 5, and F) 6 meta-bands. The upper brain plots display the network topologies used to generate the underlying meta-bands (i.e., the ground-truth network topologies), while the brain plots on the bottom row depict the network topologies of the detected meta-bands. In the central plot, the Y-axis represents the subjects, while the X-axis depicts each frequency bin. Above this plot, there is a bar indicating the frequencies where each of the original meta-bands were defined. Different colors represent different communities. The original and detected brain topologies and its corresponding meta-band are depicted with the same color. Of note, the same color can be associated to different communities across figures.*

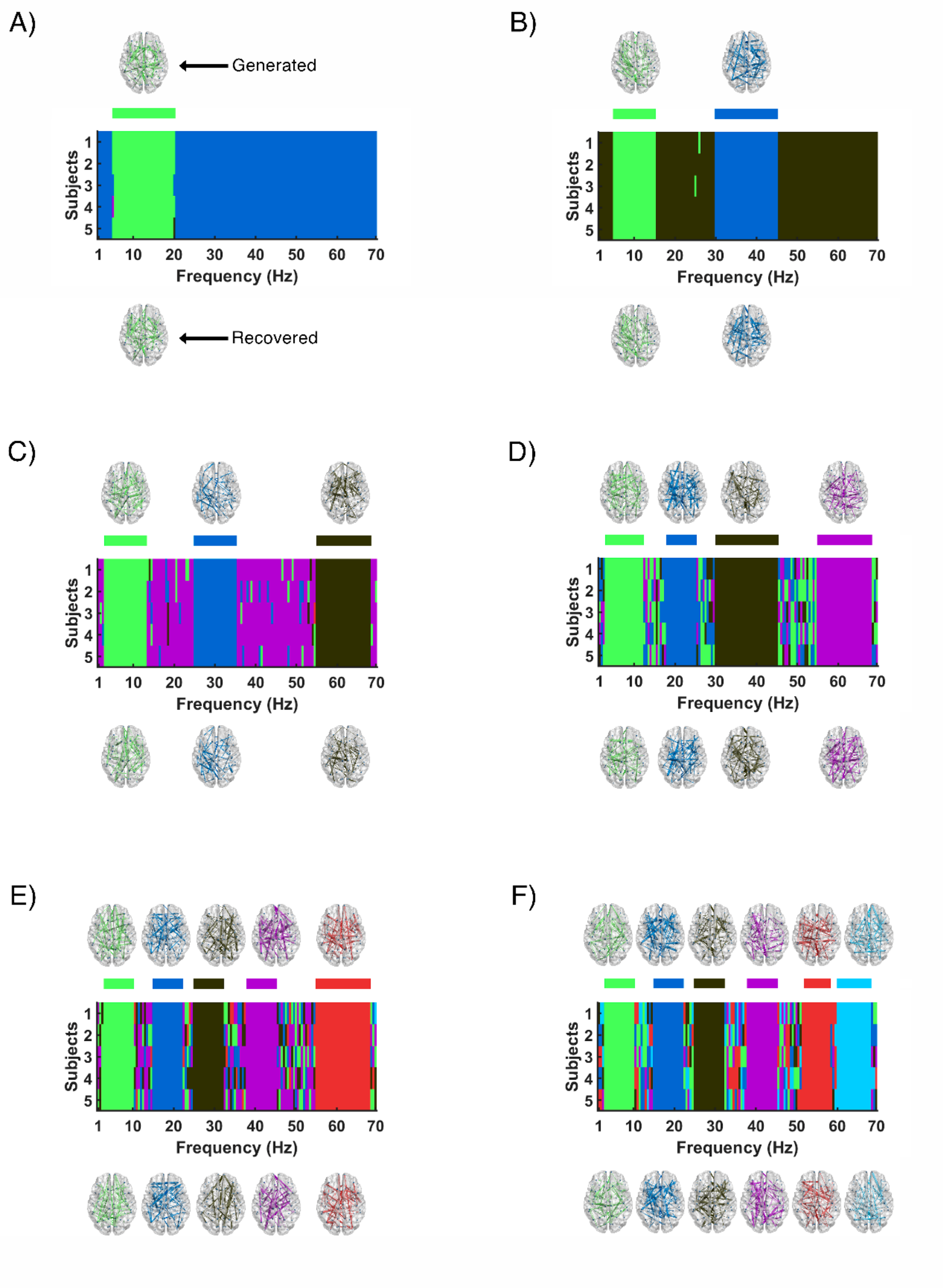

*Figure S19. Meta-bands detected with M/EEG-like synthetic signals for different numbers of underlying bands for the "Not continuous" scenario (sampling frequency = 200 Hz): A) 1, B) 2, C) 3, D) 4, E) 5, and F) 6 meta-bands. The upper brain plots display the network topologies used to generate the underlying meta-bands (i.e., the ground-truth network topologies), while the brain plots on the bottom row depict the network topologies of the detected meta-bands. In the central plot, the Y-axis represents the subjects, while the X-axis depicts each frequency bin. Above this plot, there is a bar indicating the frequencies where each of the original meta-bands were defined. Different colors represent different communities. The original and detected brain topologies and its corresponding meta-band are depicted with the same color. Of note, the same color can be associated to different communities across figures.*

### 3. ADDITIONAL ANALYSES PERFORMED WITH REAL M/EEG RECORDINGS

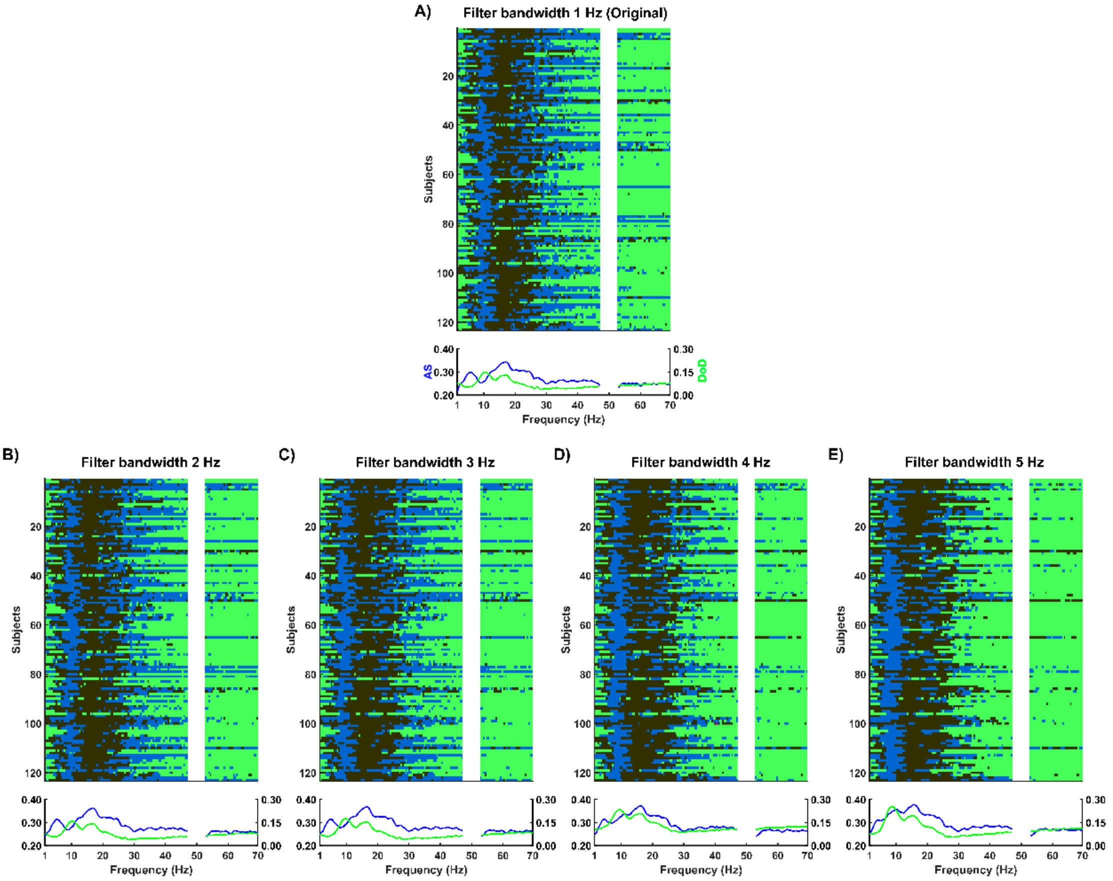

*Figure S20. Meta-band sequencing for the MEG database using a filter bandwidth of: A) 1 Hz, B) 2 Hz, C) 3 Hz, D) 4 Hz, E) 5 Hz. In the upper diagram of each panel (A-E), the Y-axis includes the Frequency Activation Sequence (FAS) for each subject, while X-axis represents each individual frequency bin, with different colors representing different communities. In the lower diagram of each panel, it is depicted the frequency evolution of the Attraction Strength (AS, blue) and the Degree of Divergence (DoD, green).*

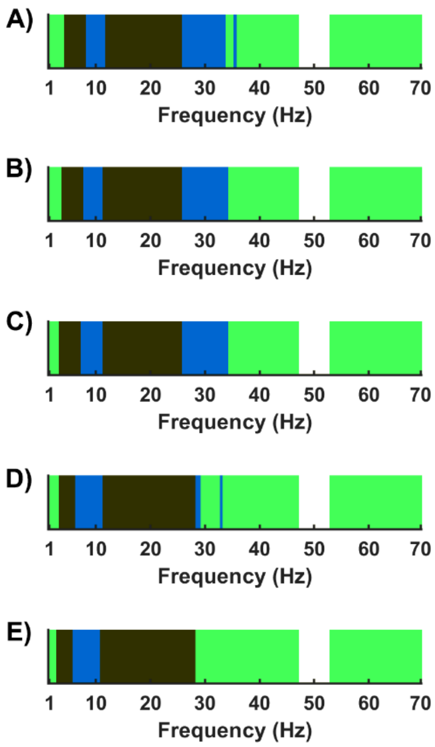

*Figure S21. Group-level mode meta-band sequencing for the MEG database when using different filter bandwidths: A) 1 Hz, B) 2 Hz, C) 3 Hz, D) 4 Hz, and E) 5 Hz.*

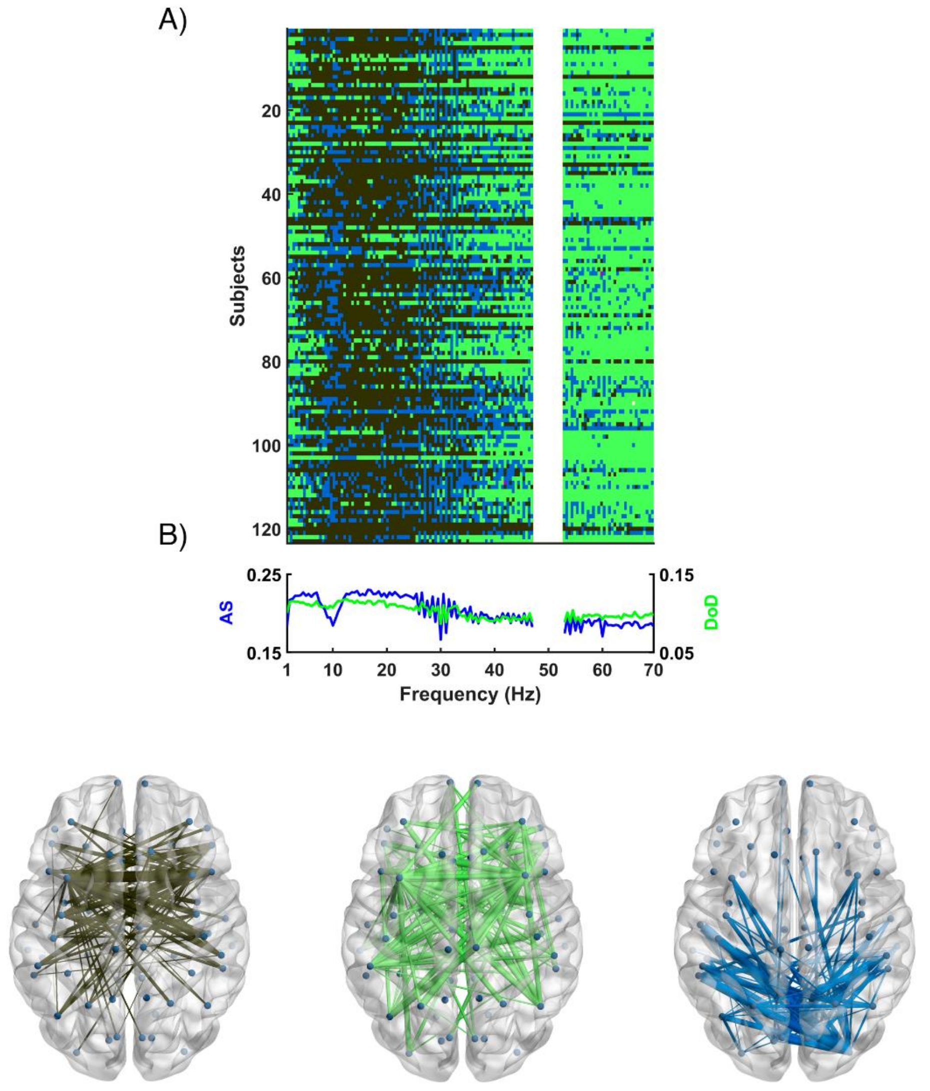

*Figure S22. Meta-bands detected with the MEG database using a filter order of 1500. A) Y-axis includes the FAS for each subject, while X-axis represents each individual frequency bin; different colors represent different communities. B) Frequency evolution of Attraction Strength (AS, blue) and Degree of Dominance (DoD, green). C) Network topologies of the three meta-bands detected; the color of each network topology corresponds with the color of the meta-bands in the upper plot.*

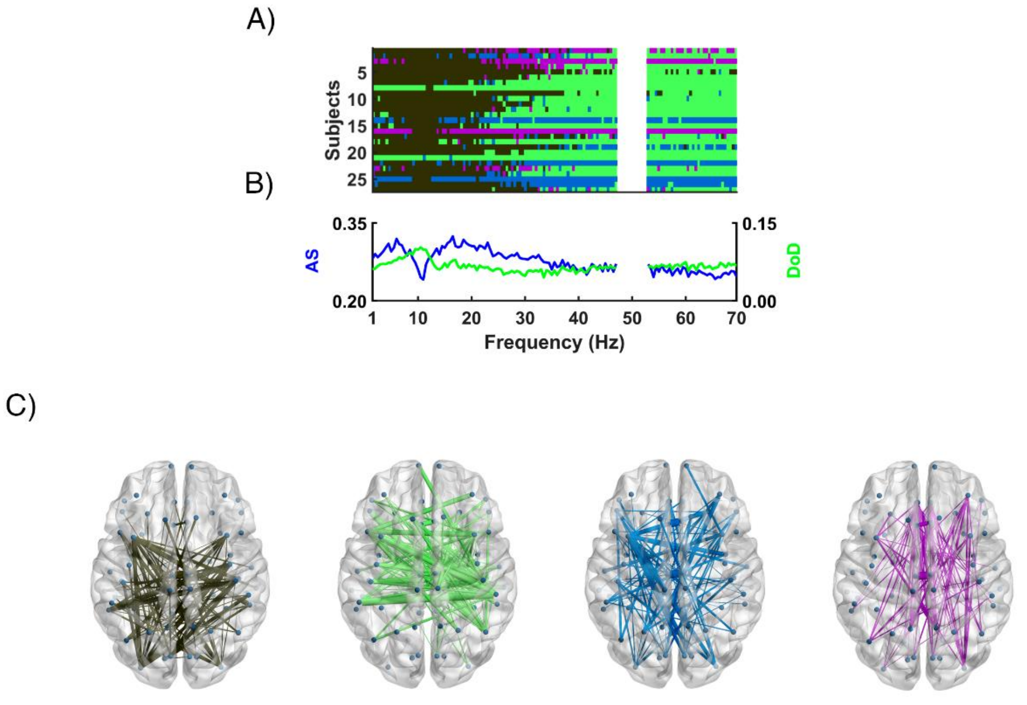

*Figure S23. Meta-bands detected with the $EEG_1$ database using a filter order of 1500. A) Y-axis includes the FAS for each subject, while X-axis represents each individual frequency bin; different colors represent different communities. B) Frequency evolution of Attraction Strength (AS, blue) and Degree of Dominance (DoD, green). C) Network topologies of the four meta-bands detected; the color of each network topology corresponds with the color of the meta-bands in the upper plot.*

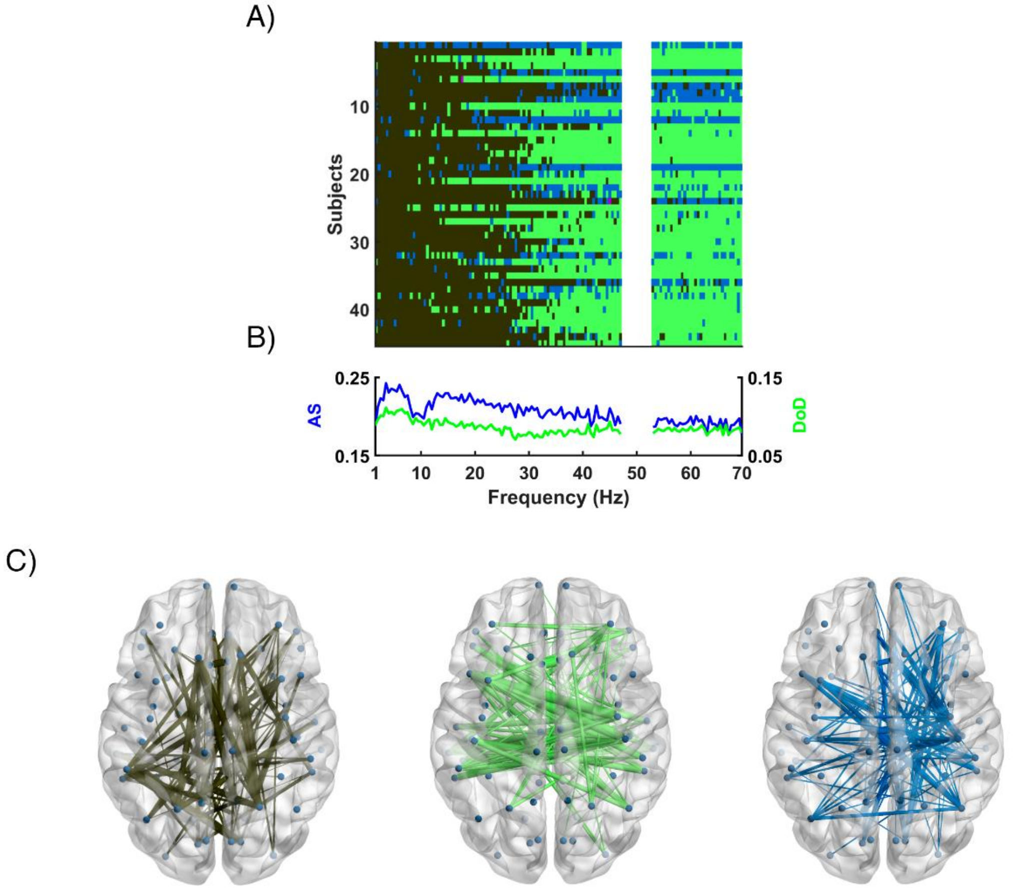

*Figure S24. Meta-bands detected with the $EEG_2$ database using a filter order of 1500. A) Y-axis includes the FAS for each subject, while X-axis represents each individual frequency bin; different colors represent different communities. B) Frequency evolution of Attraction Strength (AS, blue) and Degree of Dominance (DoD, green). C) Network topologies of the three meta-bands detected; the color of each network topology corresponds with the color of the meta-bands in the upper plot.*

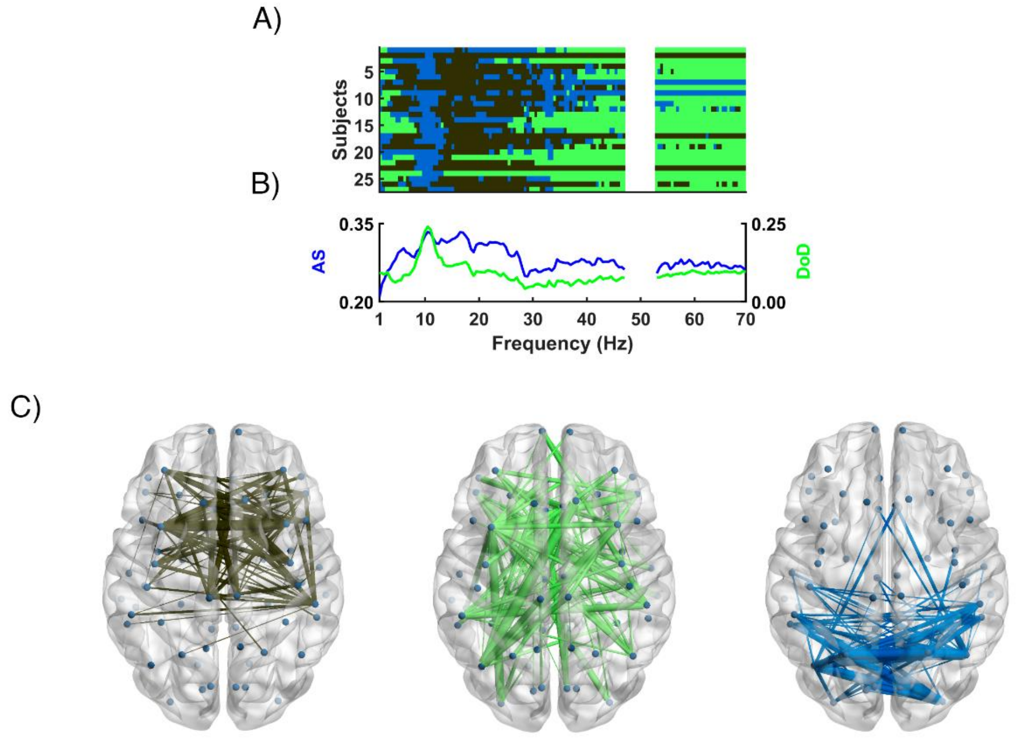

*Figure S25. Meta-bands detected with 27 random subjects of the MEG database using a filter order of 500. A) Y-axis includes the FAS for each subject, while X-axis represents each individual frequency bin; different colors represent different communities. B) Frequency evolution of Attraction Strength (AS, blue) and Degree of Dominance (DoD, green). C) Network topologies of the three meta-bands detected; the color of each network topology corresponds with the color of the meta-bands in the upper plot.*

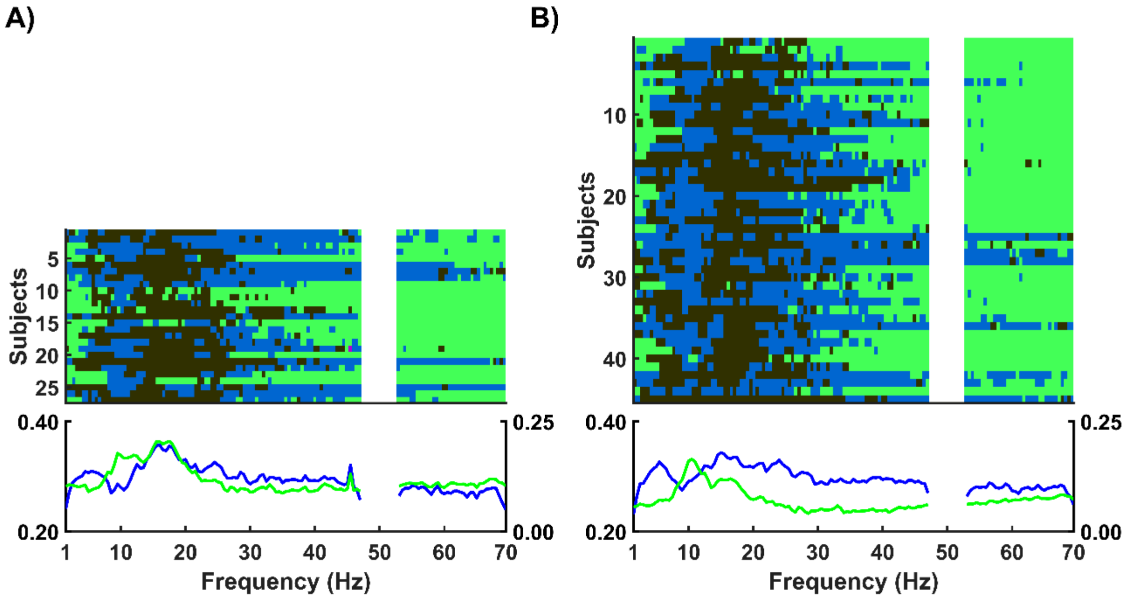

*Figure S26. Meta-band sequencing for the two new MEG subsets: A) 27 subjects with 32 coils, B) 45 subjects with 29 coils. The selected MEG coils were the closest neighbors to any of the electrodes of the corresponding EEG layout. In the upper diagram of each panel (A and B), the Y-axis includes the FAS for each subject, while X-axis represents each individual frequency bin, with different colors representing different communities. In the lower diagram of each panel, it is depicted the frequency evolution of Attraction Strength (AS, blue) and Degree of Divergence (DoD, green).*

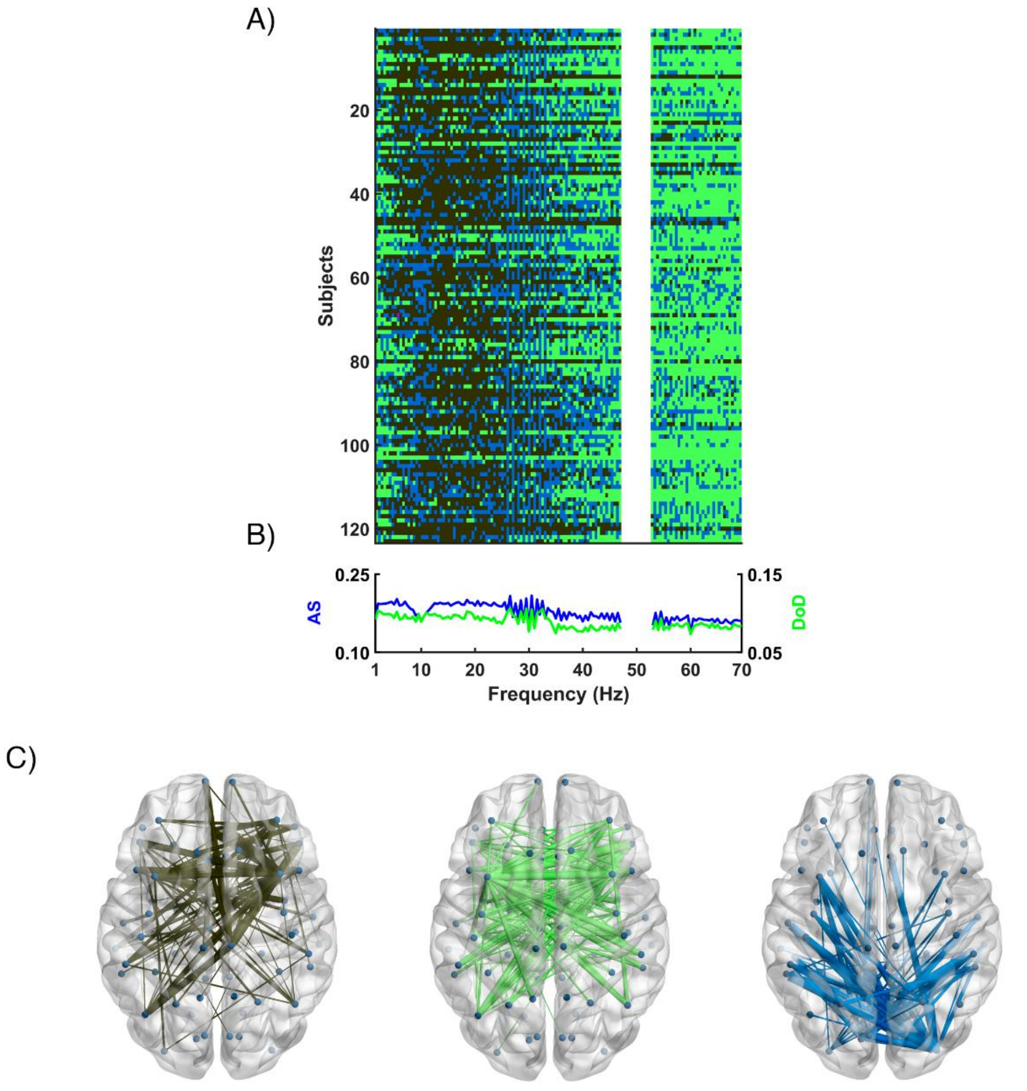

*Figure S27. Meta-bands detected with the MEG database downsampled to 200 Hz using a filter order of 500. A) Y-axis includes the FAS for each subject, while X-axis represents each individual frequency bin; different colors represent different communities. B) Frequency evolution of Attraction Strength (AS, blue) and Degree of Dominance (DoD, green). C) Network topologies of the three meta-bands detected; the color of each network topology corresponds with the color of the meta-bands in the upper plot.*
